# Anterior insular cortex plays a critical role in interoceptive attention

**DOI:** 10.1101/464867

**Authors:** Xingchao Wang, Qiong Wu, Laura Egan, Xiaosi Gu, Pinan Liu, Hong Gu, Yihong Yang, Jing Luo, Yanhong Wu, Zhixian Gao, Jin Fan

## Abstract

Although accumulating evidence indicates that the anterior insular cortex (AIC) mediates interoceptive attention, which refers the attention towards physiological signals arising from the body, the necessity of the AIC in this process has not been demonstrated. Using a novel task that directs attention toward breathing rhythm, we assessed the involvement of the AIC in interoceptive attention in healthy participants using functional magnetic resonance imaging and examined the necessity of the AIC in interoceptive attention in patients with AIC lesions. We found that interoceptive attention was associated with greater AIC activation, as well as enhanced coupling between the AIC and somatosensory area along with reduced coupling between AIC and visual sensory areas. AIC activation and connectivity were predictive of individual differences in interoceptive accuracy. Importantly, AIC lesion patients showed disrupted interoceptive discrimination accuracy and sensitivity. Together, these results provide compelling evidence that AIC plays a critical role in interoceptive attention.

## Introduction

As defined by William James in *Principles of Psychology,* attention is “taking possession by the mind, in clear and vivid form, of one out of what seem several simultaneous objects or trains of thought” (James, 1890). Thus, the target of attention can be either the external objects or the internal thoughts. Although external and internal attention have been extensively investigated, a third category of attention, the attentional mechanism in interoceptive awareness, which is the conscious focus on bodily somatic and visceral signals or responses (Craig, 2002, 2003, 2010; Critchley, 2005; Critchley, Wiens, Rotshtein, Ohman, & Dolan, 2004), i.e., the interoceptive attention, has been far less studied. Previous theories argue that subjective emotions arise from these bodily reactions and visceral experiences (Dolan, 2002) and that interoceptive awareness informs allostatic regulatory processes about the state of the body in order to maintain homeostasis (Gu & FitzGerald, 2014). Thus, appropriate attention to bodily states and accurate perception of interoceptive information are essential in emotional awareness and in the maintenance of normal physiological conditions. Recent human studies emphasize the role of the insula in interoceptive representations (Daubenmier, Sze, Kerr, Kemeny, & Mehling, 2013; Farb, Segal, & Anderson, 2013b; Ronchi et al., 2015). Neuroanatomical evidence, consistent with neuroimaging findings, suggest that the anterior insular cortex (AIC) is an important structure for encoding and representing interoceptive information (Craig, 2002, 2003, 2009; Critchley et al., 2004; Stephani, Fernandez-Baca Vaca, Maciunas, Koubeissi, & Luders, 2011).

Although the AIC has been recognized as an interoceptive cortex (Craig, 2003; Critchley et al., 2004; Ernst et al., 2014; Singer, Critchley, & Preuschoff, 2009; Terasawa, Fukushima, & Umeda, 2013), these findings remain equivocal because AIC activation seems ubiquitous across a wide range of tasks involving cognition, emotion, and other cognitive processes in addition to interoceptive attention (Gu, Hof, Friston, & Fan, 2013; Uddin, Kinnison, Pessoa, & Anderson, 2014). In addition, the correlational AIC activation found in functional neuroimaging studies alone does not provide causal evidence for its role in interoceptive attention, leaving the question of whether the AIC is critical in interoceptive attention unanswered. Studying patients with focal lesions in the AIC (e.g., in Gu et al., 2012; Gu et al., 2015; Ronchi et al., 2015; Starr et al., 2009; Wang et al., 2014) would thus provide a unique opportunity to examine the necessity of the AIC in this fundamental process.

One challenge to studying interoceptive attention is the vague nature of interoceptive awareness. With the classic definition of attention by James (1890), only the contents that are clearly perceived and represented by the mind can be the target of attention. However, most existing tasks measuring interoceptive attention fail to meet this criterion. In contrast to exteroceptive attention towards external sensory inputs, it is difficult to obtain precise measurements of interoceptive attention experimentally because of the imprecise perception of visceral changes such as heart rate (Paulus & Stein, 2010; Ring, Brener, Knapp, & Mailloux, 2015; Windmann, Schonecke, Frohlig, & Maldener, 1999). Multiple sources of physical information contribute to bodily signals and most of these sources of somatic feedback cannot be described accurately by mindful introspection in normal physiological status (Ring et al., 2015). This limitation impedes accurate measurement of interoceptive attention and examination of the neural mechanisms underlying this process. To overcome this barrier, a perceivable visceral channel needs to be used.

Breathing is an essential activity to maintain human life, and more importantly, is an easily perceivable internal bodily signal. Breathing, as an autonomous vital movement, can also be measured and actively controlled in humans (Daubenmier et al., 2013; Davenport, Chan, Zhang, & Chou, 2007). The unique physiological characteristics of respiration make breath detection an ideal method to measure interoceptive accuracy/sensitivity (Garfinkel, Seth, Barrett, Suzuki, & Critchley, 2015) and to explore the neural activity underlying interoceptive attention. Thus, we designed a breath detection task to engage interoceptive attention (attention to bodily signals), in which participants were required to indicate whether a presented breathing curve is delayed or not relative to their own breathing effort (breath detection task, BDT), in contrast to engage exteroceptive attention (attention to visual signals), in which participants were required to indicate whether a visual dot stimulus is flashed on the breathing curve (dot flash detection task, DDT). This design enabled us to examine the involvement of the AIC in interoceptive processing in healthy participants and the necessity of the AIC in this processing in patients with AIC lesions.

Based on previous evidence (e.g., Critchley, 2004), we hypothesized that AIC is critical in interoceptive attention for the integration of information from an individual’s homeostatic state and the external environment to reach subjective awareness. We first conducted fMRI studies with two samples to map the neural substrates underlying interoceptive attention to internal bodily signals in contrast to exteroceptive attention to external visual signals in healthy participants while they performed the tasks. We then investigated the necessity of AIC in interoceptive attention by assessing interoceptive attention in patients with focal AIC lesions, compared to brain-damaged controls (patients with lesions in areas other than insular or somatosensory related cortices) and matched neurologically intact controls. We predicted that the AIC would be involved in interoceptive attention and that patients with AIC lesions would show deficits in performance on the interoceptive, but not exteroceptive, attention task.

## Methods

### Task design

#### Task implementations

A respiratory transducer (TSD201, MRI compatible, BIOPAC Systems Inc.), which was fastened around the participants’ upper chest, was utilized to record breathing effort by measuring thoracic changes in circumference occurring during respiration. The signal for the change in circumference was sampled at 1000Hz using the BIOPAC MP150/RSP100C system, passing through a DC amplifier with low-pass filtering at 1Hz and high-pass filtering at .05Hz, and gain set to 10V. Analog signal was then digitized by an A/D converter (USB-1208HS-4AO, Measurement Computing, Inc.) and sent to a USB port of the test computer (Figure 1a). The task program in E-Prime^™^ (Psychology Software Tools, Pittsburgh, PA, USA) served as an interface through which the digitized signal from the USB port was received and presented to the participants on the computer screen as a continuous blue breath curve extending from left to right as time elapsed (Figure 1b), which was representative of their breathing effort. The breath curve was presented either with or without a delay (Figure 1c).

**Figure 1.**
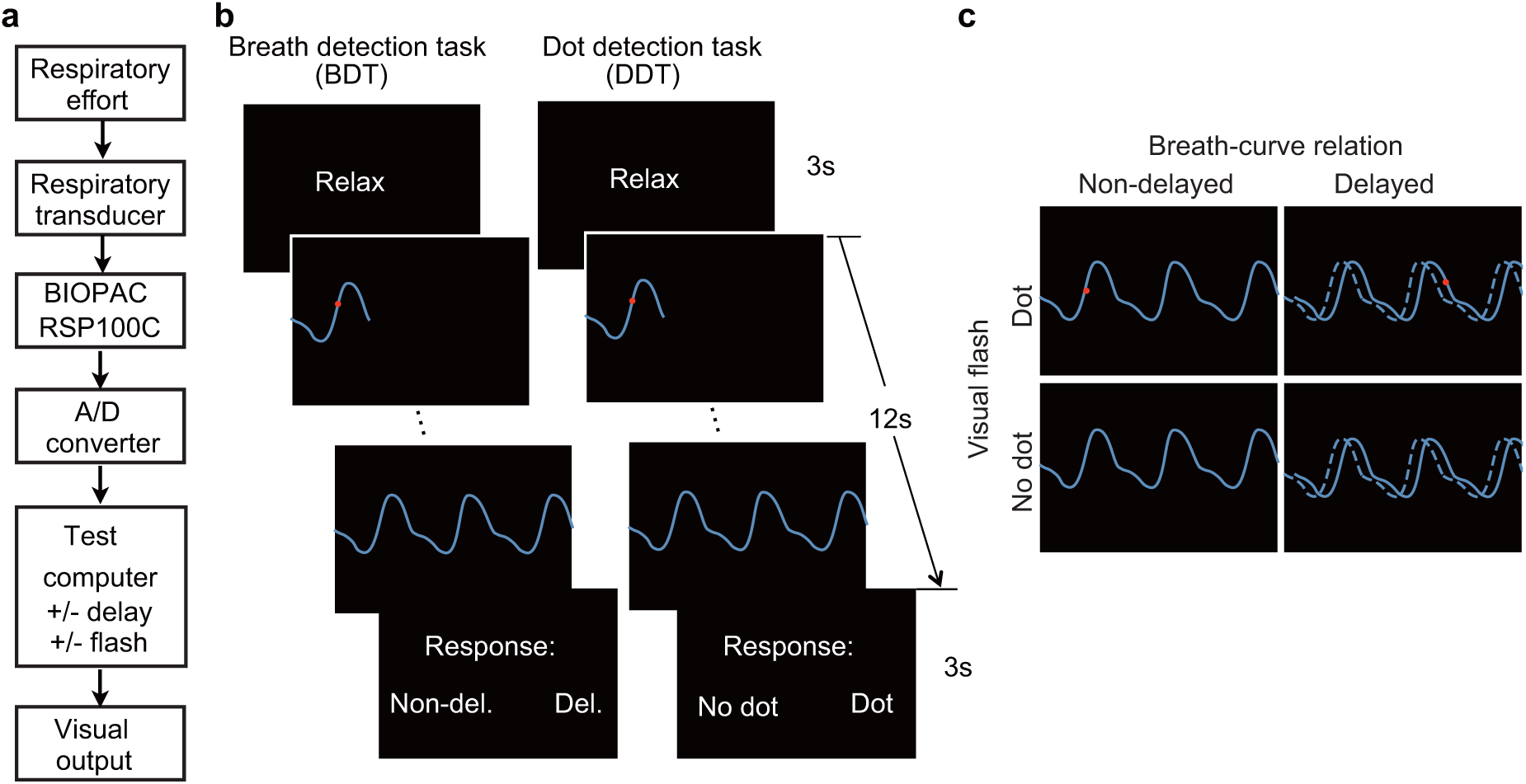
Experimental setup, trial structure of the tasks, and stimulus conditions. (a) The respiratory effort is converted to electronic signal changes using a respiratory transducer, amplified by BIOPAC, digitized using an A/D converter, and sent to the test computer for the final visual displays as a dynamic breath curve, with or without a 400-ms delay. (b) This panel shows two trials for breath delay detection task (BDT) and flash dot detection task (DDT) runs, respectively. Each trial begins with a 3-s blank display, followed by a 12-s display of respiratory curve presented with or without a 400 ms delay and with or without a 30 ms red dot flashed at a random position of the curve, and ended with a 3-s response window during which participants make a forced-choice button-press response to two alternative choices depending on the block type (BDT or DDT) to indicate whether the feedback curve was synchronous or delayed (for the BDT block) or whether a dot appeared (for the DDT block). (c) The task represents a 2 × 2 × 2 factorial design with the factors of attention to breath or dot (block design), and presence or absence of breath curve delay, and presence or absence of a dot flashed.

For the engagement of interoceptive attention during BDT, participants were required to judge whether the presented breath curve was delayed compared to the breath rhythm they perceived from their body. There were two levels for the manipulation of BDT: Non-delayed and Delayed. In half of the trials, the displayed breath curve was synchronized with the participant’s own respiration. In the other half, the displayed breath curve was delivered after a 400-ms delay period compared to the participant’s own respiration (i.e., the plotting of the point on the extending curve was actually the point saved 400 ms before the current time point). Note that the parameter of 400 ms delay was determined based on a proportion (~1/10) of an average respiratory cycle of normal healthy people which is 3~4 s/cycle. For the engagement of exteroceptive attention, the DDT was performed. Participants were instructed to detect whether a red dot flashed on the respiratory curve at any time when the breath curve was displayed. There were also two levels for the DDT: dot did not appear (No dot) and dot appeared (Dot). In half of the trials, a red dot flashed (30 ms for the fMRI experiment, and 50 ms for the lesion study) at a randomized time point on the breath curve. Figure 1b illustrates the two tasks. These two tasks constituted 4 trial conditions of a 2 (BDT: non-delayed, delayed) by 2 (DDT: no dot, dot) factorial design (Figure 1c), and therefore constituted four trial types, reflecting presence and absence of feedback delay and presence and absence of a red dot flash, presented in a random order in each run. During the two tasks, participants were instructed to breath as usual without holding or forcing their breath.

Participants were asked to perform each task in a blocked fashion in the Interoceptive and Exteroceptive runs. The fMRI experiment consisted of two runs, with one run for the interoceptive task and the other run for the exteroceptive task. There were 60 trials per run. Each run began and ended with a 30 s blank display, each trial lasted 18 s, with an average of 2 s inter trial interval, for a total of 21 minutes per run. Each trial began with a 3-s relax display, followed by a 12-s respiratory curve presented with or without an addition of 400 ms delay, and ended with a 3-s response window during which participant made a forced-choice button-press response, prompted by presentation of two alternative choices for participants to indicate their response (Figure 1b). The inter-trial interval was 2 s. After the fMRI scan, participants were asked to indicate the subjective difficulty they felt for each task on a 1 – 10 scale with higher value indicating feeling more difficult. For the lesion study, the same tasks were employed, one run for each task, with 40 trials in each run.

#### Behavioral data analysis

Interoceptive attention is associated with objective *accuracy* in detecting bodily signals, subjective belief in one’s ability to detect bodily signals in general (i.e., *sensibility*), and the correspondence between objective accuracy and subject report (i.e., metacognitive *awareness* about one’s performance when detecting bodily signals) (Garfinkel & Critchley, 2013; Garfinkel et al., 2015). Objective interoceptive/exteroceptive accuracy was calculated as the overall correct response rate during interoceptive/exteroceptive task. In addition, we also used signal detection theory to index detection sensitivity and response bias. Signal detection theory characterizes how perceivers separate signal from noise, assuming that the perceiver has a distribution of internal responses for both signal and noise (Snodgrass & Corwin, 1988; Stanislaw & Todorov, 1999). A fundamental advantage of signal detection theory is the distinction between sensitivity (ability to discriminate alternatives) and bias (propensity to categorize ‘signal’ or ‘noise’). For the BDT, the sensitivity index (*d*′) was calculated as *d*′ = *Z_hitrate_* − *Z_false alarm rate_*, where the hit rate is the proportion of trials with correct response to delayed breath curve present and response ‘yes’, and the false alarm rate is the proportion of trials with non-delayed breath curve presented and responded as ‘yes’ of delayed. A higher value of *d*′ indicates a better interoceptive sensibility, while a value of 0 represents performance at chance level. The response bias index (*β*), representing the position of the subject’s decision criterion, was defined as *β* = exp (*d*′ × *C*), where *C* = -(*Z_hit rate_* + *Z_false alarm rate_*)/2. Index *β* corresponds to the distance of participants’ estimated criterion to ideal observer criterion, and a value of 1 indicates no bias. For the DDT, indices of *d*′ and *β* were calculated using the same formula, with the dot present as ‘signal’ and dot absent as ‘noise’. Relative interoceptive accuracy was defined as the difference of performance accuracy between BDT and DDT to control for non-specific performance effects (Critchley et al., 2004).

An individual’s subjective account of how they experience internal sensation and perception represents an alternative aspect of interoceptive processing, namely sensibility (Garfinkel et al., 2015). In the fMRI experiment, the subjective sensibility of interoceptive processing was quantified using the self-report questionnaire of Body Perception Questionnaire (BPQ) (Porges, 1993). Subjective perception of one’s performance during an interoceptive task represents the awareness aspect of interoceptive attention (Garfinkel et al., 2015), which was measured via the subjectively scored difficulty of interoceptive task relative to exteroceptive task. The correlation between the relative interoceptive accuracy and these indices of subjective sensibility and awareness was calculated to examine the relationship between the perceived (subjective) and actual measured (objective) performance of interoceptive attention.

### fMRI experiments

#### Participants

fMRI experiments included two samples of participants: the first sample included 44 adults (23 females and 21 males, mean age ± standard deviation: 21.43 ± 2.51 years, age range: 19-29 years) and the second sample included additional 28 adults (14 females and 14 males, mean age ± standard deviation: 21.93 ± 2.11 years, age range: 18-26 years). All participants underwent the same experimental procedures, except that pulse and respiratory signals were recorded for the second sample using the pulse sensor (Siemens Peripheral Pulse Unit, PPU_098) of the scanner and BIOPAC, respectively. All participants (except for one participant) were right-handed, reported normal or corrected-to-normal vision, and had no known neurological or visual disorders. All participants completed questionnaires indexing subjective interoceptive sensibility (BPQ), symptoms of anxiety (Hamilton anxiety scale, HAMA (Hamilton, Schutte, & Malouff, 1976)), depression (Beck Depression Inventory, BDI (Knight, 1984)), and positive and negative affective experience (PANAS) (Watson, 1988). They gave written informed consent in accordance with the procedures and protocols approved by The Human Subjects Review Committee of Peking University.

#### fMRI data acquisition and preprocessing

During functional scanning, participants performed the interoceptive and exteroceptive tasks in separate runs that required them to attend to either their respiration or a visual flash dot, respectively. All neuroimaging data were acquired on a MAGNETOM Prisma 3T MR scanner (Siemens, Erlangen, Germany) with a 64-channal phase-array head-neck coil. During the tasks, blood oxygen level-dependent (BOLD) signals were acquired with a prototype simultaneous multi-slices echo-planar imaging (EPI) sequence (echo time, 30 ms; repetition time, 2000 ms; field of view, 224 mm x 224 mm; matrix, 112 × 112; in-plane resolution, 2 mm × 2 mm; flip angle, 90 degree; slice thickness, 2.1 mm; gap, 10%; number of slices, 64; slice orientation, transversal; bandwidth, 2126 Hz/Pixel; slice acceleration factor, 2). For the second cohort, the thickness was changed to 2 mm with the gap of 15%, and the number of slices was changed to 62. Field map images were acquired using a vendor-provided Siemens gradient echo sequence (gre field mapping: echo time 1, 4.92 ms; echo time 2, 7.38 ms; repetition time, 635 ms; flip angle, 60 degree; bandwidth, 565 Hz/Pixel) with the same geometry and orientation as the EPI image. A high-resolution 3D *T*_1_ structural image (3D magnetization-prepared rapid acquisition gradient echo; 0.5 mm × 0.5 mm × 1 mm resolution) was also acquired. Image preprocessing was performed using Statistical Parametric Mapping package (SPM12; Welcome Department of Imaging Neuroscience, London, United Kingdom). EPI volumes were realigned to the first volume, corrected for geometric distortions using the field map, coregistered to the *T_1_* image, normalized to a standard template (Montreal Neurological Institute, MNI), resampled to 2×2×2 mm^3^ voxel size, and spatially smoothed with an isotropic 8 mm full-width at half-maximum (FWHM) Gaussian kernel.

#### fMRI: whole brain analysis for the first sample

##### Image statistical parametric mapping

Imaging data from the two samples were analyzed separately and independently, with the exploratory whole brain analysis conducted with the first sample and the confirmatory region of interest (ROI) analysis conducted with the second sample. For the whole brain analysis of the first sample, statistical inference was based on a random effects approach (Penny & Holmes, 2007), which comprised two steps: first-level analyses estimating contrasts of interest for each subject followed by second-level analyses for statistical inference at the group level. For each participant, first-level statistical parametric maps of BOLD signals were modeled using general linear modeling (GLM) with regressors defined for each session with the four trial types: 2 breath curve delay (non-delayed, delayed) × 2 dot present (no dot, dot). The corresponding four regressors were generated by convolving the onset vectors of each trial type with a standard canonical hemodynamic response function (HRF). Six parameters generated during motion correction were entered as covariates of no interest. The time series for each voxel were high-pass filtered (1/128 Hz cutoff) to remove low-frequency noise and signal drift.

Contrast maps for interoceptive attention (interoceptive task – exteroceptive task), the breath curve delay (delayed – non-delayed), and the interaction between them ([delayed – non-delayed] _interoceptive task_ – [delayed – non-delayed] _exteroceptive task_) for each participant were entered into a second-level group analysis conducted with a random-effects model that accounts for inter-subject variability and permits population-based inferences. A Monte Carlo simulation using the AlphaSim program (http://afni.nimh.nih.gov/pub/dist/doc/manual/AlphaSim.pdf) was conducted to determine an appropriate cluster threshold. A cluster extent of 193 contiguous voxels was indicated as necessary to correct for multiple comparisons at *p* < 0.05 with a voxelwise *p* value at 0.001 (estimated smoothness FWHM = 11.57 mm × 12.43 mm × 11.59 mm, Resels = 208.44, Volume = 238955). Note that changes in neural activity revealed by the main effect of interoceptive attention (the contrast of interoceptive versus exteroceptive condition) could also reflect task specific effects (i.e., task difficulty) or respiratory characteristics (i.e., amplitude and frequency), in addition to effect of change in attentional focus. Although the main effect of interoceptive attention is subject to this confounding, the interaction effect should not be confounded by the change in respiratory characteristics. This interaction reflects the brain response to the mismatch (delay versus non-delayed) during interoceptive processing controlling for the non-specific effect (i.e., the difference in feedback stimulus of delay and non-delayed under the exteroception task condition). Therefore, a positive interaction effect represents brain response to the interoceptive processing above and beyond the physical feedback difference.

##### Correlation between interoceptive accuracy and the interaction effect of the AIC

To test for a linear correlation between AIC activity and behavioral performance on the interoceptive task, we entered each participant’s interaction contrast maps into the second-level random effect group regression analysis, together with their individual accuracy in the interoceptive task as the variable of interest and accuracy in the exteroceptive task as the covariate. Threshold of significance was at *p* < 0.05 with a voxelwise *p* value of 0.01 and a cluster extent of 34 contiguous voxels, corrected using small-volume region-of-interest correction for multiple comparisons using AlphaSim program. The mask image was generated from an anatomical template of bilateral insular cortex based on the Automated Anatomical Labeling (AAL) template (Tzourio-Mazoyer et al., 2002).

##### Psychophysiological interaction (PPI) analysis

PPI analysis provides a measure of change in functional connectivity between different brain regions under a specific psychological context (Friston et al., 1997). We conducted PPI analyses using a moderator derived from the product of the activity of a seed region (i.e., the AIC) and the psychological context (i.e., interoceptive in contrast to exteroceptive task). The region of interest (ROI) selection was independent of the interoceptive attention process that was used as psychological context. The left and right AIC were first identified from the main effect of the breath curve delay (the contrast of delayed versus non-delayed) in the GLM. We then conducted two whole-brain PPI tests for right and left AIC, reflecting changes in functional connectivity between the seed region time series (physiological regressor) and other brain regions as a function of interoceptive relative to exteroceptive attention (psychological regressor). The AIC time series of each participant were extracted from a 6-mm radius sphere centered at the peak of AIC (right AIC: x = 30, y = 26, z = −4; left AIC: x = −30, y = 24, z = −4). The PPI term was calculated as element-by-element product of the deconvolved physiological regressor and psychological regressor, which was then reconvolved with the canonical HRF. The PPI model generated included the PPI term, the physiological regressor, the psychological regressor, and nuisance regressors of six motion parameters. Threshold of significance for the second-level group data analysis of the images from PPI regressor was determined the same as in the GLM. Regions identified as significant clusters have two possible interpretations: (1) the connectivity between the AIC and those regions was altered by the psychological context, or (2) the response of those regions to the psychological context is modulated by AIC activity. To simplify the explanation, we used the first interpretation throughout this article.

##### Dynamic causal modeling (DCM)

DCM (Friston, Harrison, & Penny, 2003) is a Bayesian model comparison procedure to infer effective connectivity between brain regions by estimating direct interactions on neuronal level. DCM distinguishes between endogenous coupling and context-specific coupling, which could account for the effects of experimentally controlled network perturbations. Due to the inherent limited causal interpretability of the PPI analysis for the direction of interaction, we only conducted DCM to explain the potential mechanisms of the interplay between AIC and other brain areas involved during interoceptive attention. The ROI of the right AIC in the DCM was the same as in the PPI analysis. The other regions included in the DCM were selected based on significant positive and negative PPI results and with the coordinates of the ROIs identified by the group level T-contrast of all conditions versus baseline. Data from six participants was excluded from DCM analysis because activity in one of the ROIs could not be identified.

A three-area DCM was specified for all participants with bidirectional endogenous connection between right AIC and the two other ROIs, and with the main effect of “all stimuli” as the driving input entering into the two other ROIs. Five base models were generated by specifying possible modulations of interoceptive and exteroceptive attention on the four endogenous connections between ROIs. These base models were then systematically elaborated to produce 52 variant models, which include all possible combination of the modulation of interoceptive and exteroceptive attention on endogenous connections between right AIC and the two other ROIs.

Model comparison was implemented using random-effects (RFX) Bayesian model selection (BMS) in DCM12 to determine the most likely model of the 52 models given the observed data from all participants (Stephan, Penny, Daunizeau, Moran, & Friston, 2009). The RFX analysis computes exceedance and posterior probabilities at the group level, and the exceedance probability of a given model denotes the probability that this model is more likely than all other models considered (Stephan et al., 2009). To summarize the strength of effective connectivity and its modulation quantitatively, we used random effects Bayesian model averaging (BMA) to obtain average connectivity estimates (weighted by their posterior model) across all models and all participants (Penny et al., 2010). We conducted one-sample t tests on the subject specific BMA parameter estimates to assess their consistency across subjects with Bonferroni correction for multiple comparisons.

#### fMRI: ROI analyses for the second sample

Whereas whole brain analyses of the first sample aimed at identifying brain areas involved in interoceptive processing, ROI analyses of the second sample aimed to confirm that the effects found from the first sample were not confounded with the effect induced by other physiological signals. Change in BOLD signals can be due to both direct neural activity (induced by experimental manipulation) and indirect effect (such as vascular response, which would be considered as a confounding effect). For example, the cerebral vascular response is sensitive to circulation of CO_2_ and O_2_, and causes a change in global cerebral blood flow (CBF) and global BOLD signal. It is evident both in human and animals that the global CBF and global BOLD responses influence local stimulus-induced hemodynamic response to neural activation (Cohen, Ugurbil, & Kim, 2002; Friston et al., 1990; Ramsay et al., 1993). Typically, a larger local stimulus-induced BOLD response occurs when global BOLD is lowered, while a smaller local stimulus-induced BOLD response occurs when global BOLD is elevated. In our study, the experimental manipulation of interoception was likely to cause a change in respiratory characteristics (i.e., circulation of CO_2_ and O_2_). The difference in physiologic states between the interoceptive and exteroceptive task conditions might cause a change in global BOLD signals, and thus the effect resulting from local interoception-related BOLD response would be confounded by the global hemodynamic influence.

To partial out the potential confounding, physiological data, including cardiac pulsation and respiratory volume collected in the second sample, were processed using PhLEM toolbox (http://sites.google.com/site/phlemtoolbox/). Physiological noise correction was implemented by regressing out cardiac- and respiratory-related effects, and their interaction effect, as noise from the fMRI signal using a modification of conventional RETROICOR approach (Brooks et al., 2008; Glover, Li, & Ress, 2000). In RETROICOR, a cardiac phase calculated from a pulse oximeter was assigned to each acquired image in a time series (Hu, Le, Parrish, & Erhard, 1995), and a respiratory phase was assigned to a corresponding image using the histogram equalized transfer function that takes both the respiratory timing and depth of breathing into account (Glover et al., 2000). Conventional RETROICOR approach (Glover et al., 2000) defines low-order Fourier terms (i.e., sine and cosine values of the principal frequency and the 2^nd^ harmonic) to model the independent effects of the cardiac and respiratory fluctuation, which is considered insufficient to remove variations caused by physiological artifacts (Harley & Bielajew, 1992; Tijssen, Jenkinson, Brooks, Jezzard, & Miller, 2014). Therefore, we used additional terms of higher-order Fourier expansions (i.e., to the 5th harmonics) in RETROICOR, and formed multiplicative sine/cosine terms that take the interaction between cardiac and respiratory effect into account. Specifically, the interaction terms were calculated by **sin(*φ_c_* + *φ_r_*)** and **cos(*φ_c_* + *φ_r_*)** where ***φ_c,r_*** is the cardiac or respiratory phase, consisting of a mixture of 3^rd^ order cardiac and 2^nd^ order respiratory harmonics. Therefore, the modified RETROICOR approach contained a total of 44 regressors, with 20 of them from independent cardiac and respiratory effects and 24 of them from interaction terms. Statistical parametric maps were generated using the same GLM as in the above whole brain analyses, with motion parameters and these physiological regressors entered as covariates of no interest.

To avoid double dipping, ROIs were defined based on the first sample. Specifically, ROIs (i.e. left and right AIC) were the clusters of the second-level group analysis results of the interaction effect ([delayed – non-delayed] _interoceptive task_ – [delayed – non-delayed] _exteroceptive task_). Parameter estimates were extracted from each ROI in the second sample for each of the 4 experimental conditions of each participant, and then entered into a two-way repeated measures analysis of variance model. We also examined correlations between the interaction effect of each ROI and behavior measures (i.e., relative interoceptive accuracy) across participants.

To further confirm that the result was not dependent on the (independent of) ROI selection, we explored the influence of physiological noise by comparing the whole brain activations related to interoception without and with physiological correction at an extremely permissive threshold (voxelwise *p* < 0.05 uncorrected).

### Lesion study

#### Brain lesion patient and control groups

Six male patients (33-53 years old, mean 42.17 ± 7.31 years) with focal unilateral insular cortex lesions participated in the lesion study (see Supplementary Table 1 for patient characteristics). Two patients had a right-side lesion, and four patients had a left-side lesion. In addition, six patients with focal lesions in regions other than insular cortex (i.e., temporal pole, n = 3, lateral frontal cortex, n = 2, and superior temporal gyrus, n = 1) were recruited as brain-damaged controls (BDCs), and 12 neurologically intact participants were recruited as normal controls (NCs). All lesions resulted from surgical removal of low-grade gliomas. All patients were recruited from the Patient’s Registry of Tiantan Hospital, Beijing, China. NC participants were recruited in the local community. All NC participants were right-handed, had normal color vision, and reported no previous or current neurological or psychiatric disorders. BDC patients matched with patients with insular cortex lesion in chronicity, and neither group significantly differed from NC group in age and education (*p*s > 0.05). All six insular lesion patients were considered cognitively intact, measured by Mini-Mental State Examination (MMSE), a measurement of cognitive impairment (Folstein, Folstein, & McHugh, 1975), and the raw scores of MMSE were not statistically different from either BDCs (*p* > 0.05) or NCs (*p* > 0.05). Patients with insular lesions did not show alteration in baseline mood as indexed by BDI score, compared to NCs or BDCs (*p*s >0.05). See Supplementary Table 1 for demographic information of the groups. Note that by chance, all of the AIC lesion patients were male. We conducted additional statistical analyses to compare the AIC patients to only male controls to examine whether there was the potential confound of gender difference (see lesion study results). All participants were informed of the study requirements and provided written consent prior to participation. The patient study was approved by the Institutional Review Board of the Beijing Tiantan Hospital, Capital Medical University.

#### Lesion reconstruction

Two neurosurgeons, blind to the experimental design and behavioral results, identified and mapped the lesions of each patient onto a template derived from a digital MRI volume of a normal control (ch2bet.nii) embedded in the MRIcro program (http://www.cabiatl.com/mricro/mricro/index.html). In each case, lesions evident on MRI were transcribed onto corresponding sections of the template in order to create a volume of interest image. This was used to measure the location (in MNI coordinates) and volume (in ml) of individual lesions and to create within group overlaps of lesions using the MRIcro program.

#### Behavioral data analysis of the lesion study

We used the non-parametric bootstrapping method (Hasson, Avidan, Deouell, Bentin, & Malach, 2003; Mooney & Duval, 1993) to assess the probability of observing a significant difference between two groups (AIC versus NC, AIC versus BDC) because the small sample data sets did not meet the assumption of parametric tests. The bootstrapping procedure was conducted with 10,000 iterations (details referred to (Gu et al., 2012)). If the probability of obtaining the observed t-value is less than 5% (one-tailed), we considered the difference between the two groups to be significant. We used one-tailed tests because of the hypothesis that lesions of a specific brain region (i.e., the AIC) would induce deficits in behavioral response. In addition, we calculated Bayes factors ∣ uniform (B_u_) to determine the relative strength of evidence for null and alternative hypotheses (Dienes, 2014; Dienes & McLatchie, 2018). The value of B means that the data are B times more likely under the alternative than under the null hypothesis. The conventional standard for assessing substantial evidence for the null is a value of B < 1/3 and for the theory against null is a value of B > 3, while values between 1/3 and 3 are counted as data insensitivity. The Bayes factors were calculated using free online Bayes calculator (http://www.lifesci.sussex.ac.uk/home/Zoltan_Dienes/inference/Bayes.htm) by specifying a uniform distribution with all population parameter values from the lower to the upper limit equally plausible.

## Results

### Behavioral results of the fMRI studies

Performance accuracy (%) and discrimination sensitivity (*d*′) in the interoceptive task were 82.1 ± 14.7% and 2.2 ± 1.1 (Mean ± SD) for the first sample, and 74.9 ± 9.6% and 1.6 ± 0.6 (Mean ± SD) for the second sample, which were significantly above chance level (50% and 0 for accuracy and *d*′ respectively; For the first sample: *t*(43) = 14.51, *P* < 0.001, Cohen’s *d* = 2.18 for accuracy and *t*(43) = 13.09, *p* < 0.001, Cohen’s *d* = 2.0 for *d*′, respectively; For the second sample: *t*(27) = 13.77, *p* < 0.001, Cohen’s *d* = 2.59 for accuracy and *t*(27) = 12.89, *p* < 0.001, Cohen’s *d* = 2.67, respectively), but lower than exteroceptive task accuracy of 87.3 ± 9.8% and *d*′ of 2.6 ± 0.8 for the first sample (*t*(43) = −2.36, *p* = 0.02, Cohen’s *d* = 0.35 and *t*(43) = −2.31, *p* = 0.03, Cohen’s *d* = 0.35, respectively), and accuracy of 80.9 ± 14.7% and *d*′ of 2.2 ± 1.1 for the second sample (*t*(27) = −1.83, *p* = 0.08, Cohen’s *d* = 0.35 and *t*(27) = −2.83, *p* = 0.009, Cohen’s *d* = 0.50, respectively). Participants were slower in terms of reaction time (RT) (only for the first sample) and less biased in interoceptive task than in the exteroceptive task (RT: *t*(43) = 2.89, *p* = 0.006, Cohen’s *d* = 0.44 for the first sample, and *t*(27) = 0.6, *p* = 0.55, Cohen’s *d* = 0.12 for the second sample; β: *t*(43) = −2.62, *p* = 0.01, Cohen’s *d* = 0.39 for the first sample, and *t*(27) = −4.32, *p* < 0.001, Cohen’s *d* = 0.80 for the second sample) (see Figure 1-figure supplement 1 and 2 for details of the behavior results for the first and the second sample respectively, i.e., accuracy, RT, *d*′, and β). Across the two samples, the split-half reliability of the interoceptive and exteroceptive tasks were 0.86 and 0.85 for the first sample, and 0.68 and 0.89 for the second sample, respectively, reflecting a high internal consistency reliability of each of these two tasks.

For the first sample, the relative interoceptive accuracy was positively correlated with the “awareness of bodily processes” subtest of body perception questionnaire (Pearson r = 0.29, p < 0.05, one-tailed) and negatively correlated with the subjectively scored difficulty of interoceptive task relative to exteroceptive task (Pearson r = −0.43, p < 0.01, one-tailed), demonstrating the validity of the measure of interoceptive accuracy. In addition, relative interoceptive accuracy was positively correlated with trait positive affective experience (measured by Positive and Negative Affect Schedule, PANAS) (Pearson r = 0.31, p < 0.05, one-tailed), suggesting a close relationship between explicit interoceptive attention towards bodily signals and subjective emotion experiences. No significant correlations were observed between relative interoceptive accuracy and anxiety (Pearson r = −0.007, p = 0.96) or depression score (Pearson r = −0.002, p = 0.99). For the second sample, however, we did not find significant correlations between relative interoceptive accuracy and scores of questionnaires (awareness of bodily processes: Pearson r = −0.17, p = 0.40; trait positive affective experience: Pearson r = 0.12, p = 0.56; anxiety: Pearson r = 0.29, p = 0.14; depression: Pearson r = 0.03, p = 0.89; note that we did not collect subjective score of task difficulty in the second sample).

### Imaging results of the whole brain analysis of the fMRI study of the first sample

#### Main effects of interoceptive attention and feedback delay, and their interaction

Interoceptive attention, compared to exteroceptive attention, was associated with enhanced activity in the cognitive control network (Fan, 2014; Q. Wu et al., 2015; Xuan et al., 2016), including bilateral AIC, dorsal ACC and supplementary motor area (SMA), and superior frontal and parietal cortices (frontal eye field, FEF; and areas near/along intraparietal sulcus, IPS; Figure 2a, Supplementary Table 2). In addition, this contrast revealed significant less activation, or deactivation, in the core regions of the default mode network (Raichle et al., 2001), including ventral medial prefrontal cortex (vmPFC), middle temporal gyrus, and posterior cingulate cortex (PCC).

**Figure 2.**
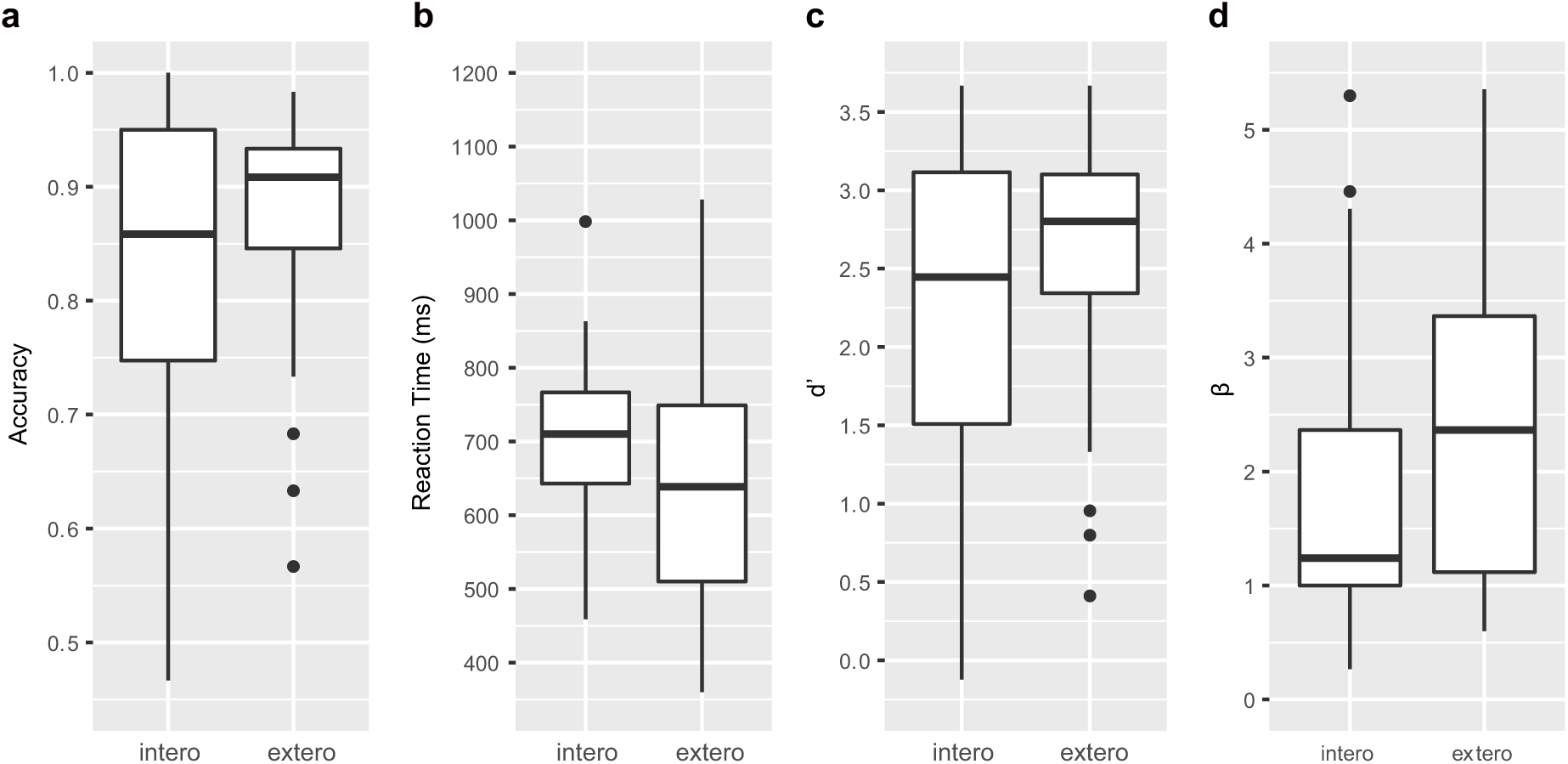
Main effects and the interaction for the whole brain analysis of the first sample. (a) Main effect of interoceptive attention (interoceptive task vs. exteroceptive task). (b) Main effect of breath curve feedback condition (delayed curve vs. non-delayed curve). (c) Interaction between attention type and breath-curve feedback condition ([delayed – non-delayed]_interoceptive task_ – [delayed – non-delayed]_exteroceptive task_). (d) The activation of the left and right AIC activity under the four task conditions, and the pattern of the interaction.

Activation in the AIC, middle frontal gyrus (MFG), SMA, and temporal parietal junction (TPJ) was associated with the main effect of feedback delay, with delayed trials inducing greater activity than non-delayed trials (Figure 2b, Supplementary Table 3). The regions showing the main effect of feedback delay also showed the interaction effect between attentional focus (interoceptive and exteroceptive) and feedback (with and without delay) (Figure 2c, Supplementary Table 4). The task-induced responses extracted from bilateral AIC, defined by the attention by feedback interaction map, shows the activation pattern under different task conditions (Figure 2d). Specifically, (1) bilateral AIC demonstrated higher activity during the interoceptive task than during the exteroceptive task, irrespective of the feedback type; (2) the mismatched delayed trials induced greater activation in bilateral AIC in comparison to non-delayed trials only during the interoceptive task. The evidence of this interaction effect in the AIC suggests that the AIC was actively engaged in interoceptive processing.

#### Correlation between interoceptive accuracy and AIC activity

Voxel-wise regression analysis revealed the relationship between the interoceptive task-induced activity strength (map of the interaction contrast) and participants’ interoceptive accuracy (performance accuracy in the interoceptive task), with exteroceptive accuracy (performance accuracy in the exteroceptive task) controlled as a covariate. The greater interaction effect of bilateral AIC (and middle temporal gyrus, MTG) was associated with higher interoceptive accuracy across participants (Figure 3a, Supplementary Table 5). The AIC activation during the interoceptive task involved attending to physiological signals and matching bodily signals to external visual input, which predicts individual differences in interoceptive attention (see Figure 3b for the illustration of the effects).

**Figure 3.**
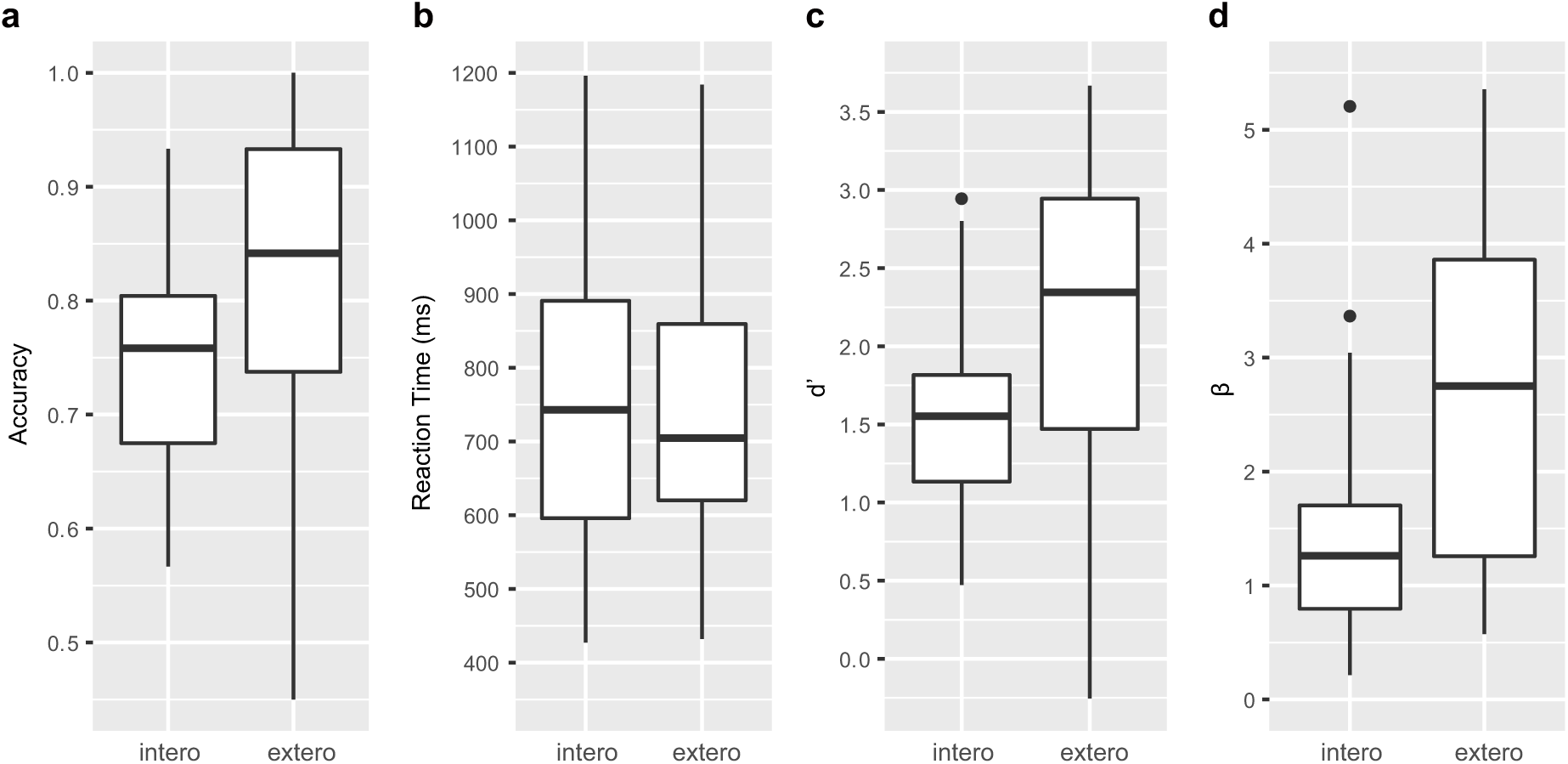
The relationship between brain activity and behavioral performance across participants. (a) This was revealed in a regression analysis of contrast images for the interaction between interoceptive attention deployment (interoception vs. exteroception) and breath curve feedback condition (delayed vs. no-delayed), with performance accuracy on interoceptive and on exteroceptive tasks as regressor-of-interest and covariate, respectively. AIC, anterior insular cortex; MTQ middle temporal gyrus. (b) The correlational patterns between the interaction effect of bilateral AIC and relative interoceptive accuracy. The data were normalized as z-scores.

#### Functional and effective connectivity of the AIC

PPI analysis showed augmented connectivity between the right AIC (as the seed) and SMA/ACC, FEF, inferior frontal gyrus (IFG), and the postcentral gyrus (PoCG) during interoceptive (versus exteroceptive) attention, in contrast to reduced connectivity between the right AIC and visual area modulated by interoceptive attention (Figure 4a, Supplementary Table 6), indicating that an increase in activity in the right AIC was associated with a greater increase in activity in FEF, IFG, and the PoCG, and more decreased activity in visual area under the condition of interoceptive compared to exteroceptive attention (Figure 4b). We found similar PPI results when using the left AIC as the seed (Figure 4-figure supplement 1).

**Figure 4.**
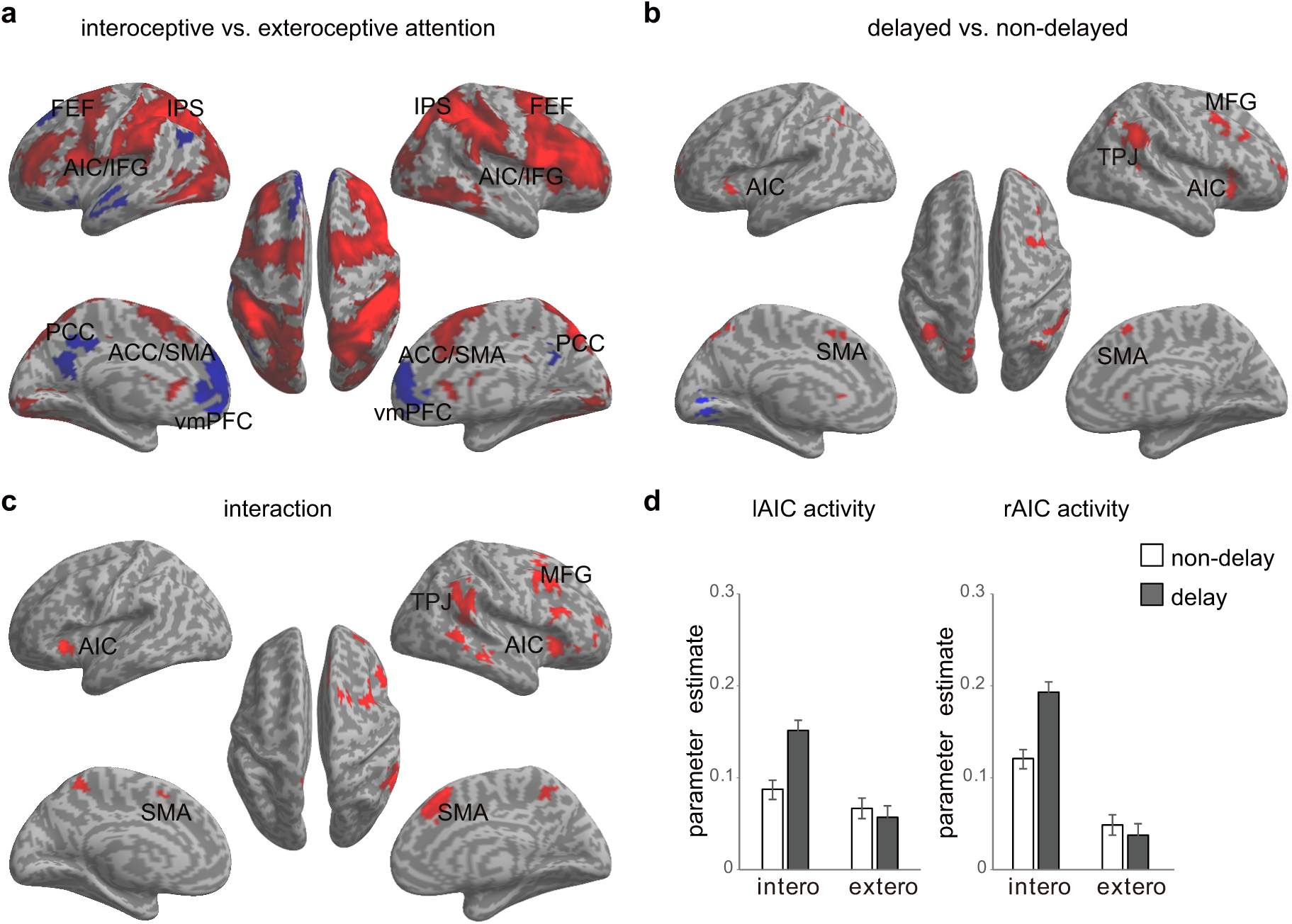
PPI and DCM results of the first fMRI sample. (a) Regions showing positive (red) and negative (blue) associations with AIC activation modulated by interoceptive attention (relative to exteroceptive attention). (b) An increase in activation in the right AIC was associated with an increase in activation in the posterior central gyrus (PoCG) and a decrease in activation in visual cortex (VC, V2/3) under the condition of interoceptive attention. (c) Five base models generated by specifying possible modulations of interoceptive and exteroceptive attention on the four endogenous connections between ROIs. The model with a dashed-line rectangle surrounded indicates the winning model revealed by fixed-effects and random-effects Bayesian model selection (BMS). (d) The intrinsic efferent connection from the AIC to the PoCG was significant. The modulatory effect of interoceptive attention on the connection from the AIC to the PoCG was significant. The modulatory effect of exteroceptive attention on the connection from AIC to V2/3 was significant (uncorrected).

Based on the PPI results, visual cortices (VC) of right V2/3 (x = 14, y = −90, z = 28 as indicated by negative PPI) and the right PoCG (x = 58, y = −16, z = 32 as indicated by positive PPI) were included in the DCM model. Note that data from one participants was excluded because significant activation in V2/3 region of interest could not be identified. For model comparison, random-effects (RFX) BMS indicated that the winning model (with an exceedance probability of 29.84%) was the one with the modulatory effects of interoceptive and exteroceptive attention exerting on the connection from the AIC to the PoCG and from the AIC to V2/3 (Figure 4c and Figure 4-figure supplement 2). The BMS indicated that interoceptive and exteroceptive attention are achieved through modulating the top-down connectivity from AIC to those two sensory cortices.

We performed parameter inference by using BMA, which takes uncertainty into account by pooling information across all models in a weighted fashion (Stephan et al., 2010). For BMA (Figure 4d), the modulatory effect of interoceptive attention was significant on the connection from the AIC to the PoCG (*t*(42) = 4.85, Bonferroni corrected *p* < 0.001). The modulatory effect of exteroceptive attention on the connection from the AIC to V2/3 was significant without correction (*t*(42) = 2.25, uncorrected *p* = 0.03). The BMA results were consistent with the winning model selected by model comparison and also PPI results: the modulatory effect from the AIC to the PoCG was driven by interoceptive attention, while the modulatory effect from the AIC to V2/3 was driven by exteroceptive attention. In addition, the BMA results highlighted the importance of the intrinsic efferent connection from the AIC to the PoCG in the network (*t*(42) = 3.61, Bonferroni corrected *p* = 0.01).

### ROI analysis results of the fMRI study of the second sample

The interaction between attentional focus (interoceptive and exteroceptive) and feedback (with and without delay) was significant in both left and right AIC (left: *F*(1,27) = 5.77, *p* = 0.024; right: *F*(1,27) = 5.73, *p* = 0.024; Figure 5a), which confirmed the interaction effect in the bilateral AIC revealed by whole brain analyses found in the first sample. The main effect of attentional focus was significant in right AIC with higher activity during the interoceptive task than during the exteroceptive task (*F*(1,27) = 9.11, *p* = 0.005), but not significant in left AIC (*F*(1,27) = 2.87, *p* = 0.10). The main effect of the feedback was not significant in neither left nor right AIC (left: *F*(1,27) < 1, *p* = 0.46; right: *F*(1,27) < 1, *p* = 0.45). In addition, we found a significant correlation between the interaction effect of right AIC and relative interoceptive accuracy (Pearson *r* = 0.36, *p* = 0.03, one-tailed), while the correlation between the interaction effect of left AIC and relative interoceptive accuracy was marginally significant (Pearson *r* = 0.29, *p* = 0.06, one-tailed; Figure 5b). To further illustrate that the interaction effect of AIC is not subject to breathing effort difference, we showed the pattern of the respiratory volume under different experimental conditions (see Figure 5-figure supplement 1). Although there was a change in the respiratory volume between interoceptive and exteroceptive task conditions (*F*(1,27) = 15.88, *p* < 0.001), this difference was canceled out by using the interaction effect (*F* < 1).

**Figure 5.**
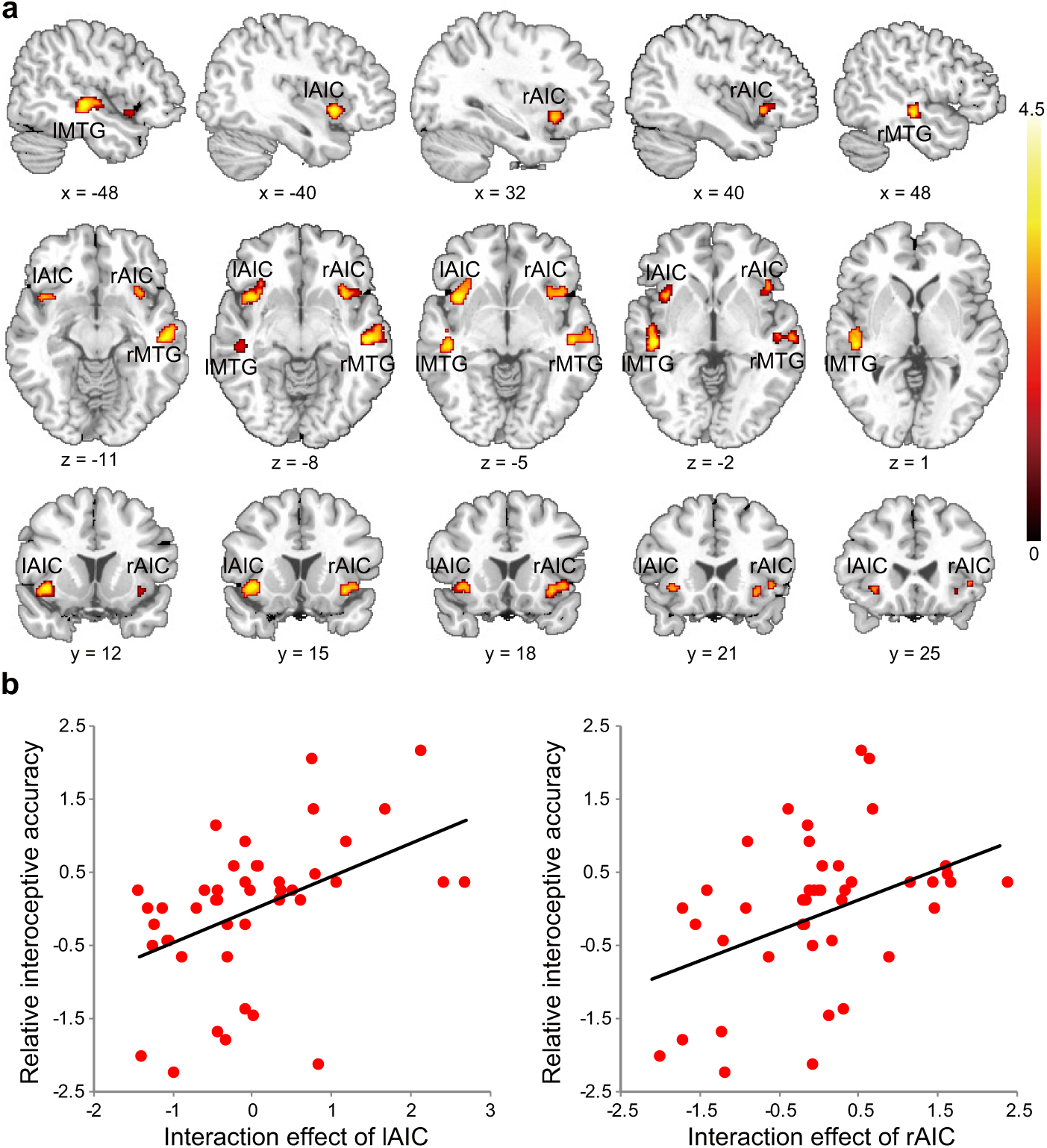
fMRI ROI results of the second sample. (a) ROI analysis of the parameter estimates of the left and right AIC for 4 experimental conditions. Error bars represent 95% confidence intervals. (b) The correlation between the interaction effect of bilateral AIC and relative interoceptive accuracy. The data were normalized as z-scores.

The whole brain analysis of the second sample showed significant overlap in the main and the interaction effects between without and with physiological correction (see Figure 5-figure supplement 2). We further checked that the AIC ROI results were not dependent on the (independent) ROI selection by comparing the whole brain activation without and with physiological correction at an extremely permissive threshold (*p* < 0.05 uncorrected). The difference of the signals of the AIC between the analyses with and without physiological corrections was not significant, suggesting that the effect of the AIC was not significantly impacted by the physiological noises (see Figure 5-figure supplement 3). Altogether, these ROI results confirmed the fact that AIC was actively engaged in interoceptive processing.

### Lesion study results: the necessity of the AIC in interoceptive attention

Figure 6 shows the insular lesion overlap for the AIC patient group. The area with the most overlap was identified as the AIC according to the literature (Kurth, Zilles, Fox, Laird, & Eickhoff, 2010; Naidich et al., 2004). For the BDT, patients with AIC lesions had significantly lower performance accuracy (58%, *t*(13) = −3.47, *p* < 0.001, B_u[0,1.0]_ = 32.94, Cohen’s *d* = 1.92 compared to NC; *t*(8) = −2.35, *p* = 0.009, B_u[0,1.0]_ = 4.11, Cohen’s *d* = 1.66 compared to BDC) (Figure 6a) and discrimination sensitivity (*d*′) compared to the NCs and BDCs groups (*t*(13) = 3.62, *p* < 0.001, B_u[0,4.0]_ = 53.32, Cohen’s *d* = 2.00 compared to NC; *t*(8) = −2.22, *p* = 0.013, B_u[0,4.0]_ = 3.56, Cohen’s *d* = 1.60 compared to BDC) (Figure 6b), indicating diminished interoceptive attention, while we did not find significant difference between NC and BDC (accuracy: *t*(8) = 0, *p* = 0.3, B_u[0,1.0]_ = 0.09; *d*′: *t*(8) = 0.112, *p* = 0.23, B_u[0,4.0]_ = 0.16). Patients with AIC lesions did not show significant alteration in propensity to categorize ‘signal’ or ‘noise’ (*β*) during interoceptive task (i.e., delayed vs. non-delayed) (AIC vs. NC: *t*(16) = 1.08, *p* = 0.09, B_u[0,4.0]_ = 0.32; AIC vs. BDC: *t*(7) = 0.37, *p* = 0.18, B_u[0,4.0]_ = 0.09; BDC vs. NC: *t*(13) = −1.51, *p* = 0.044, B_u[0,4.0]_ = 0.45) (Figure 6c). For the DDT, in contrast, patients with AIC lesions did not show abnormalities in performance, as measured by accuracy (AIC vs. NC: *t*(9) = 0.18, *p* = 0.22, B_u[0,1.0]_ = 0.09; AIC vs. BDC: *t*(7) = −0.99, *p* = 0.10, B_u[0,1.0]_ = 0.17; BDC vs. NC: *t*(16) = 1.74, *p* = 0.03, B_u[0,1.0]_ = 0.29), *d*′ (AIC vs. NC: *t*(9) = 0.18, *p* = 0.22, B_u[0,4.0]_ = 0.14; AIC vs. BDC: *t*(7) = −0.83, *p* = 0.12, B_u[0,4.0]_ = 0.26; BDC vs. NC: *t*(16) = 1.46, *p* = 0.05, B_u[0,4.0]_ = 0.39), and *β* (AIC vs. NC: *t*(15) = −0.47, *p* = 0.17, B_u[0,4.0]_ = 0.23; AIC vs. BDC: *t*(10) = −0.11, *p* = 0.23, B_u[0,4.0]_ = 0.14; BDC vs. NC: *t*(14) = −0.34, *p* = 0.19, B_u[0,4.0]_ = 0.21) compared to the NC and BDC groups (Figure 6d-f). Our results demonstrate significant impairment in discrimination ability when attending to bodily signals, but not to external visual input, in patients with AIC lesion.

**Figure 6.**
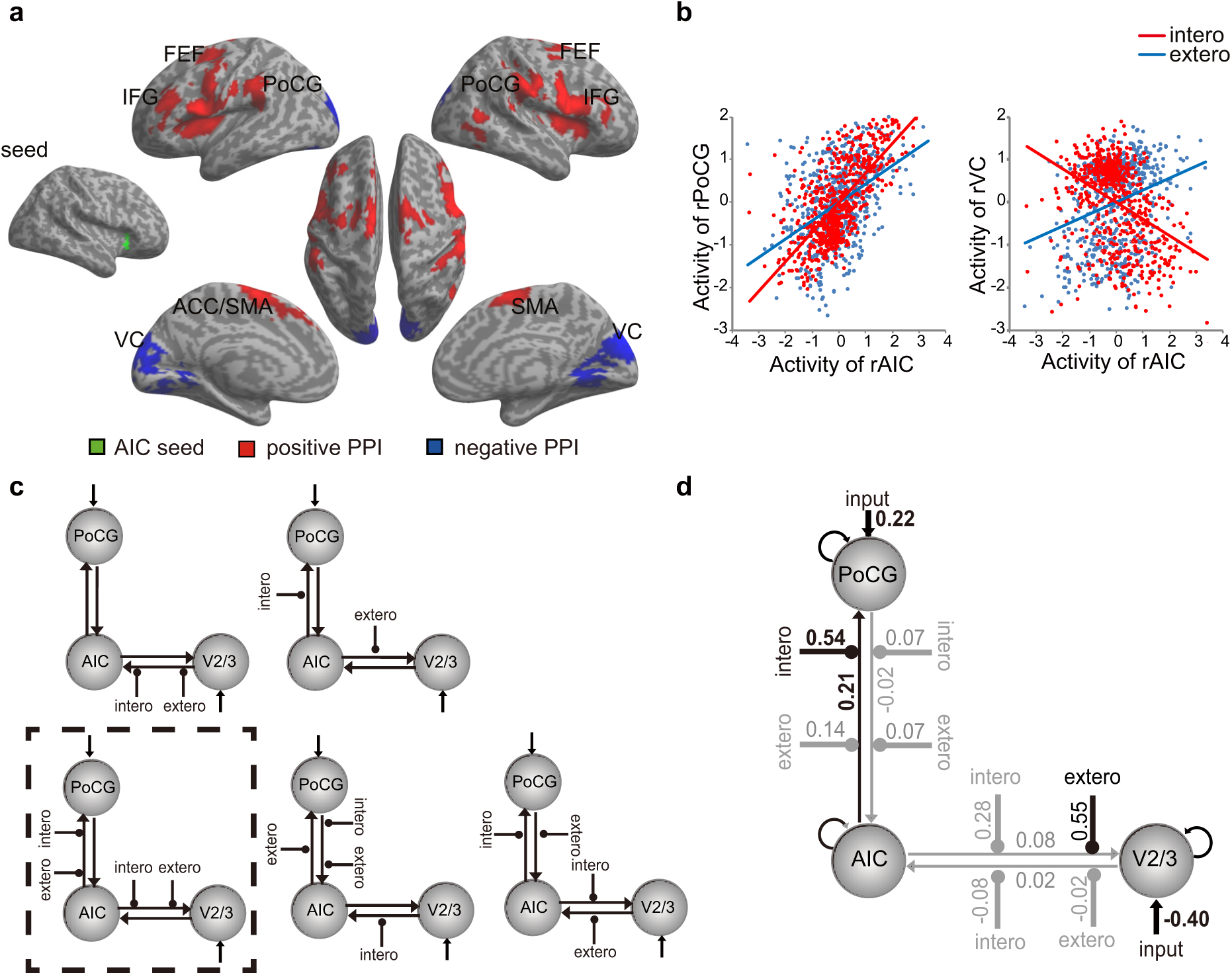
Reconstruction of anterior insular cortex lesions of six patients. Red color indicates 100% overlap. Left lesions were flipped to the right side in order to map the lesion overlap.

**Figure 7.**
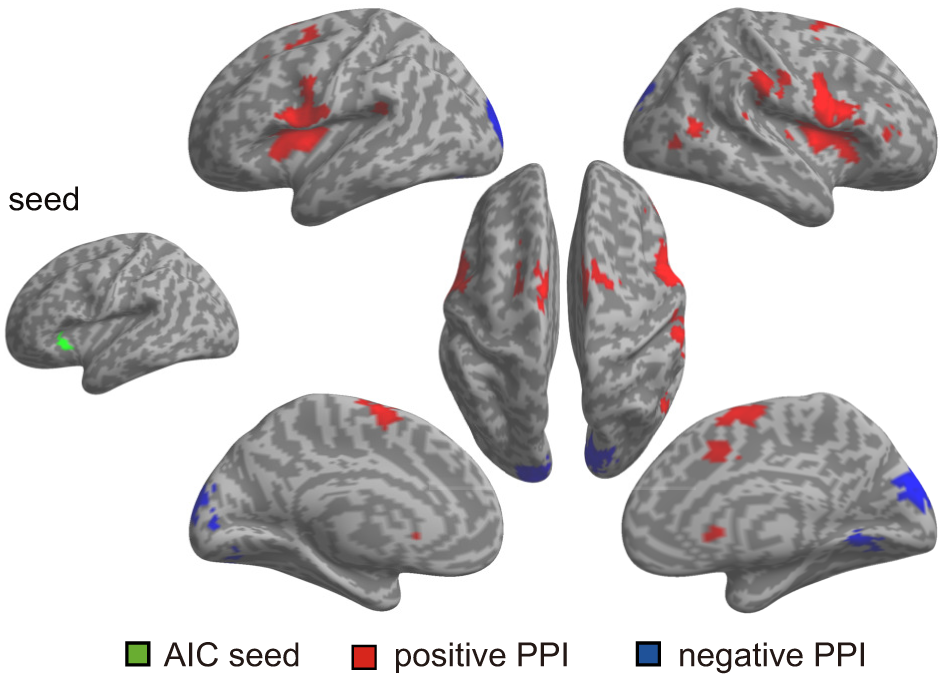
Behavioral results of the lesion study. (a, b, c) the interoceptive task performance, and (d, e, f) the exteroceptive task performance. For the interoceptive attention task, patients with AIC lesions had significantly lower performance in accuracy and *d*′ compared to NC and BDC groups, but did not show significant alteration in *β* during interoceptive task. For the exteroceptive task, patients with AIC lesions did not show abnormality in performance in accuracy, *d*′, and *β* compared to either the NC or BDC groups. NC, normal control; BDC, brain damage control. Dashed line: chance level.

To test the influence of gender on task performance during interoceptive and exteroceptive attention, we compared performance discrimination sensitivity *d*′ between males and females in NC and BDC. Neither group showed significant gender difference on both measures in the interoceptive and exteroceptive tasks (*ps* > 0.05). However, male patients with AIC lesions had worse performance accuracy and *d*′ than the NC (accuracy: *p* < 0.001; **d*′: p* < 0.001) and BDC (accuracy: *p* < 0.01; **d*′: p* = 0.01) groups. These results provided additional evidence that deficits in interoceptive attention were associated with AIC lesions without significant gender influence and support the argument that the results were not biased by the fact that all of the AIC lesion patients were male.

## Discussion

Using fMRI, we showed that the AIC is involved in interoceptive attention towards respiration supported by the underlying connectivity between the AIC and somatosensory cortex and visual areas for regulating interoceptive and exteroceptive attention. In addition, self-report of positive affective experience was positively correlated with interoceptive accuracy (though not significant in the second fMRI sample), which supports the notion that interoceptive attention is critical in emotional awareness (Barrett, Quigley, Bliss-Moreau, & Aronson, 2004; Bechara & Naqvi, 2004; Caruana, Jezzini, Sbriscia-Fioretti, Rizzolatti, & Gallese, 2011; Critchley et al., 2004; Gu et al., 2013; Pollatos, Kirsch, & Schandry, 2005; Seth, 2013; Terasawa, Moriguchi, Tochizawa, & Umeda, 2014; Wiens, 2005). Notably, we confirmed the necessity of the AIC in supporting interoceptive attention by showing reduced behavioral performance on the interoceptive task in patients with focal AIC lesions. Thus, this study demonstrates that the AIC plays a critical role in interoceptive attention.

### The necessity of the AIC for interoceptive attention

Previous functional neuroimaging studies have shown that the insula is activated in autonomic arousal and emotional reactions (Craig, 2002, 2003; Critchley et al., 2004) and emphasized the central role of the insula in interoceptive awareness. The achievement of interoceptive awareness depends on the integration of afferent bodily signals with higher order contextual information attributable to the AIC (Craig, 2002, 2009; Critchley, 2005; A. R. Damasio et al., 2000; Mutschler et al., 2009). In this study, an increase in neural activation in the AIC and other related brain structures when focusing on breath rhythm indicates that the AIC supports attention toward bodily signals. Most importantly, participants’ performance accuracy on the interoceptive task was significantly correlated to the activity of the AIC, further demonstrating the importance of the AIC in interoceptive attention.

Anatomically, the bilateral insula receives thalamo-insular projections of the interoceptive pathways (Craig, 2002). The AIC encodes subjective feelings (Craig, 2003, 2009; Flynn, Benson, & Ardila, 1999) and is critical for instantaneous representation of the state of the body (Gu & FitzGerald, 2014; Gu et al., 2015). During the BDT, this is achieved by attending to bodily signals (i.e., breath rhythm) and matching them to the external visual feedback (i.e., breath curve). The present results provide additional support to previous findings that the activity of the right AIC is related to accuracy in sensing the timing of one’s bodily signals, e.g., heartbeats (Critchley et al., 2004). Consistent with the notion that the AIC contributes to accurate perception of bodily states (Bechara & Naqvi, 2004), the insula works as a hub to convey bodily information into internal feelings for maintaining homeostasis and to mediate representations of visceral states that link to representations of the external world (Farb, Segal, & Anderson, 2013a).

The critical role of the AIC in interoceptive attention identified by the fMRI data was augmented by the data from patients with focal insula damage. Relative to non-insular lesion patients and healthy controls, AIC damage led to a deficit in accuracy and sensitivity of interoceptive attention. These findings provide causal evidence demonstrating the critical role of the AIC in interoceptive attention. Traditionally, the insular cortex is considered to be a limbic sensory region that participates in the intuitive processing of complex situations (Augustine, 1996; Butti & Hof, 2010; Menon & Uddin, 2010) by integrating ascending visceromotor, somatosensory inputs with attention systems via intrinsic connectivity to identify and respond to salient stimuli (Menon & Uddin, 2010; Uddin, 2015). The AIC, in particular, is a node mediating cognitive processes including bottom-up control of attention (Corbetta, Kincade, & Shulman, 2002; Corbetta, Patel, & Shulman, 2008) and conscious detection of signals arising from the autonomic nervous system (Craig, 2002; Critchley, 2004). Therefore, the behavioral deficit of interoceptive attention in AIC lesion patients is due to the disruption in the integration of the somatic and visceral inputs with the abstract representation of the present internal state (i.e. the saliency of certain type of signals). Consequently, it leads to the failure in discriminating whether the displayed respiratory curve is different from internal states.

### Mechanisms of the AIC in relation to interoceptive attention

We can view interoceptive attention as the mechanism that coordinates the processing of bodily signals and higher-level representation of that information. It has been proposed that the AIC encodes and represents bodily information (e.g., visceral states) and transmits this information to other neural systems for advanced computations in conscious perception and decision-making (Bechara & Naqvi, 2004; Flynn et al., 1999; Gu & FitzGerald, 2014). The AIC is a key node of the large-scale network that detects information from multiple sources including objective visceral signals and generates subjective awareness (Craig, 2009; Gu et al., 2012; Gu et al., 2015; Kleckner et al., 2017; Seeley et al., 2007), as well as responding to the switch between networks that supports internal oriented processing and cognitive control (Menon, 2011; Menon & Uddin, 2010; Sridharan, Levitin, & Menon, 2008). Supporting this argument, we showed that the AIC is intrinsically connected to the somatosensory area of the PoCG and that this connection is positively modulated by interoceptive attention (relative to exteroceptive attention).

Other higher-level areas, e.g., the ACC/SMA, FEF, and IFG of the so-called cognitive control network (CCN) (Fan, 2014; T. Wu et al., 2017), are also involved in the interoceptive process. This is also supported by the results of the enhanced functional connectivity between the AIC and these regions. Both somatosensory afferents and a network that includes the AIC and ACC are possible pathways of interoceptive attention (Khalsa, Rudrauf, Feinstein, & Tranel, 2009; Rudrauf et al., 2009). The AIC may play a central role in integrating sensory signals from the PoCG and visual cortex and send top-down signals that guide sensation and perception through a dynamic interaction with sensory or bottom-up information. Somatosensory information concerning the internal state of the body is conveyed through the PoCG, as well as the visual signals in V2/3 containing the majority of external information. The top-down modulation of AIC in interoceptive attention is accomplished by augmenting the efferent signals to the somatosensory cortices. This is consistent with the argument that a first-order mapping of internal feeling is supported by insular and somatosensory cortices (A. Damasio, 2003) and that somatosensory information critically contributes to interoceptive attention (Khalsa et al., 2009).

In the BDT, interoceptive attention reflects combination of attention to the internal bodily signal (i.e., the breath) and the external visual stimulus (i.e., the curve). In order to better coordinate perceptual processing, the AIC may distribute and balance the processes of external and internal information. The winning model and parameter inference from DCM provides evidence that interoceptive attention is achieved mainly by modulating the connectivity between AIC and somatosensory areas (PoCG), while exteroceptive attention is primarily modulated via the connectivity between the AIC and V2/3. We propose that the dynamic adjustment of the connectivity of the AIC to sensory cortices is the foundation of interoceptive attention for bodily signals, which is critical for homeostatic regulation, and of exteroceptive attention for external objects or inputs.

### The interoceptive task in the respiratory domain

Although the neural correlates of interoceptive awareness have been studied by other tasks such as heartbeat detection task (Bechara & Naqvi, 2004; Critchley et al., 2004; Khalsa et al., 2009; Ring et al., 2015), the error rate arising from the difficulty in heartbeat counting or non-sensory process confounds are inherent to these cardioception designs (Kleckner, Wormwood, Simmons, Barrett, & Quigley, 2015; Ring et al., 2015). In contrast to cardioception, breath can be clearly perceived and autonomously controlled.

This feature made the present study on interoceptive attention, which requires that the target of interoception be clearly and vividly perceivable by our consciousness. The positive correlations between objective interoceptive accuracy during the BDT and subjectively scored difficulty of interoceptive task relative to exteroceptive task (i.e., interoceptive awareness) further demonstrate that the BDT is valid in assessing interoceptive attention. We developed the BDT as a non-intrusive measurement with low cognitive load that is more practical for patients with focal brain damage than the demanding cardioception tasks.

It should be noted that the BDT does not represent a pure probe of interoception because respiratory processes can also be tracked using exteroceptive and proprioceptive information. It thus seems likely that participants relied on a mix of interoceptive, exteroceptive, and proprioceptive information to perform the task. In our design, we included the DDT for a measure of exteroception so that the subtraction of BDT and DDT leaves the interoceptive and proprioceptive processing components of interoception, which are the two out of three components of exteroceptive, interoceptive, and proprioceptive systems (Gu et al., 2013). In the BDT, the delayed manipulation in our study was fixed to 400 ms, approximately 1/10 of an average cycle of normal healthy people which is 3~4 s/cycle. It is worth trying to manipulate this delayed duration according to each individual’s respiratory cycle in an effort to control subjective task difficulty across participants.

### Interoceptive attention

Depending on the source of information, attention can be categorized into (1) interoceptive attention, which is directed toward bodily signals such as somatic and visceral signals (e.g., in a heartbeat detection or counting task); (2) exteroceptive attention, which is directed toward primary sensory inputs from outside (e.g., visual and auditory stimuli); and (3) executive control of attention, which coordinates thoughts and actions (e.g., in Color Stroop, flanker, and working memory tasks; see review Fan (2014)). Although there are extensive studies on the attentional modulation of sensory and perceptual inputs and on the executive control of attention, it is difficult to study interoceptive attention because the vast majority of intrinsic visceral activity, except breath effort, cannot be clearly perceived under normal conditions. Using the BDT to examine attentional deployment toward breath effort enabled us to reveal the neural mechanism of interoceptive attention. In general, the perceptible, controllable, measurable, and autonomous features of breathing guaranteed more accurate and reliable measurement of individual differences in interoceptive ability.

As a kind of interoceptive attention that could be clearly perceived and autonomously controlled by people, the breath plays potentially important role in generating and regulating emotion. For example, mindfulness meditation, which is now well known for its role in emotion regulation and mental health (Khoury, Sharma, Rush, & Fournier, 2015), can be viewed as a practice involving interoceptive attention. One of its primary methods is to bring one’s attention (the processing) and then awareness (the outcome) to the current experiences of the movement of the abdomen when breathing in and out or the breath as it goes in and out the nostrils. Given the revealed neural mechanisms of interoceptive attention in this study, it would be intuitive to predict that the AIC play an important role in meditation. Findings that meditation experience is associated with increased gray matter thickness in the AIC (Lazar et al., 2005) and increased gyrification (increase in folding) of the AIC (Luders et al., 2012) support this prediction. Meditation training may enhance interoceptive attention to focus on the bodily signals so that the mind can be released from the intensive involvement of exteroceptive (the external attention) and executive control of attention (the internal attention for the coordination of thought processing) that consume the majority of mental resources.

## Conclusion

In summary, we provided important evidence of the involvement of the AIC in interoceptive attention by our fMRI study and further demonstrated that the AIC is critical by the lesion study. The converging evidence from these studies also suggests that interoceptive attention is achieved through top-down modulation from the AIC to the somatosensory and sensory cortices. In addition, the implementation of the interoceptive task extends the research on interoceptive processing into the respiratory domain with the validity and reliability demonstrated. It may have significant applications in studying issues related to interoceptive attention in patients with neuropsychiatric disorders such as anxiety (Avery et al., 2014) and autism (Barrett & Simmons, 2015; Quattrocki & Friston, 2014) and in patients with substance use disorders (Sönmez, Kahyacı Kılıç, Ateş Çöl, Görgülü, & Köse Çımar, 2017).

## Acknowledgements

We thank Dr. Thomas Beck and Dr. Tian-Yi Qian from Siemens Healthcare for providing the simultaneous mullti-slices EPI sequence for fMRI data acquisition. This work was supported by the National Natural Science Foundation of China (grant number: 81729001, 81328008) to J.F. & Z.G.; J.F. was also supported by the National Institute of Mental Health (NIMH) of the National Institutes of Health (NIH) under Award Number R01 MH094305. Y.W. was supported by the research grant of 973 (grant number: 973-2015CB351800) and National Natural Science Foundation of China (grant number: 31771205, 61690205). Y.Y. and H.G. were supported by the Intramural Research Program, National Institute on Drug Abuse, NIH. X.W. and P.L. were supported by Beijing Municipal Science & Technology Commission (grant number: Z161100002616014). X.W. was also supported by National Natural Science Foundation of China (grant number: 81600931), Beijing Municipal Administration of Hospital’ Youth Programs (code: QML20170503) and Capital Health Development Research Project of Beijing, China (grant number: 2016-4-1074). Q.W. was supported by China Postdoctoral Science Foundation (grant number: 2016M600835). X.G. was supported by the Dallas Foundation and a faculty startup grant from the University of Texas at Dallas. Dr. Nicholas Van Dam and Evelyn Ramirez were involved at the early stage of the study on interoception.

## Figure Captions

**Figure 1-figure supplement 1.**
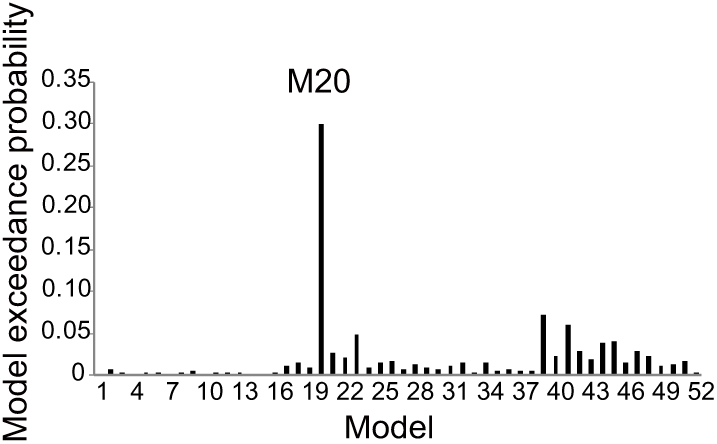
Box plots visualizing the five-number summary (minimum, lower quartile, median, upper quartile and maximum) for (a) accuracy, (b) reaction time, (c) ď, and (d) beta for BDT and DDT tasks in the first sample of fMRI study. The dots are outliers.

**Figure 1-figure supplement 2.**
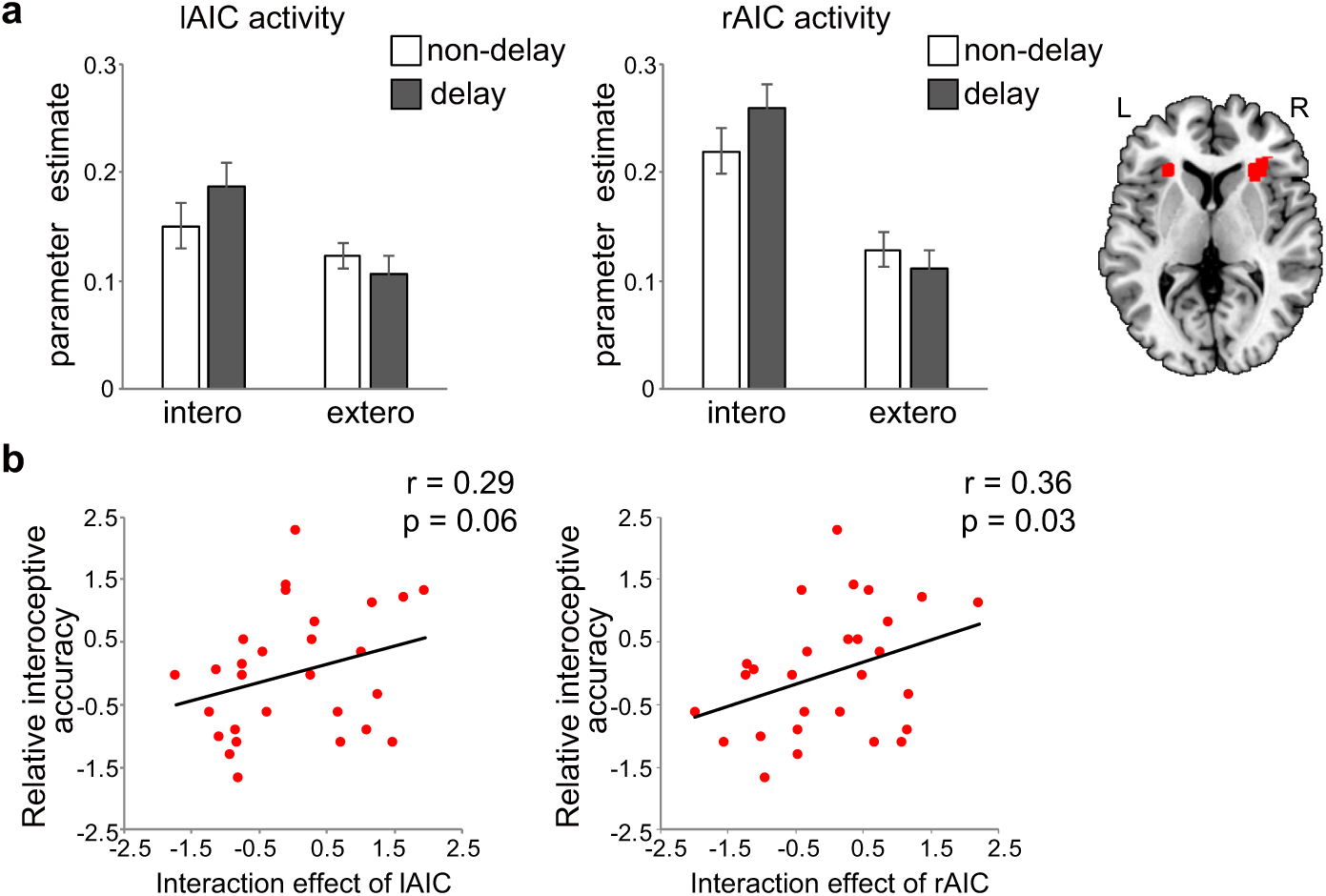
Box plots visualizing the five-number summary (minimum, lower quartile, median, upper quartile and maximum) for (a) accuracy, (b) reaction time, (c) ď, and (d) beta for BDT and DDT tasks in the second sample of fMRI study. The dots are outliers.

**Figure 4-figure supplement 1.**
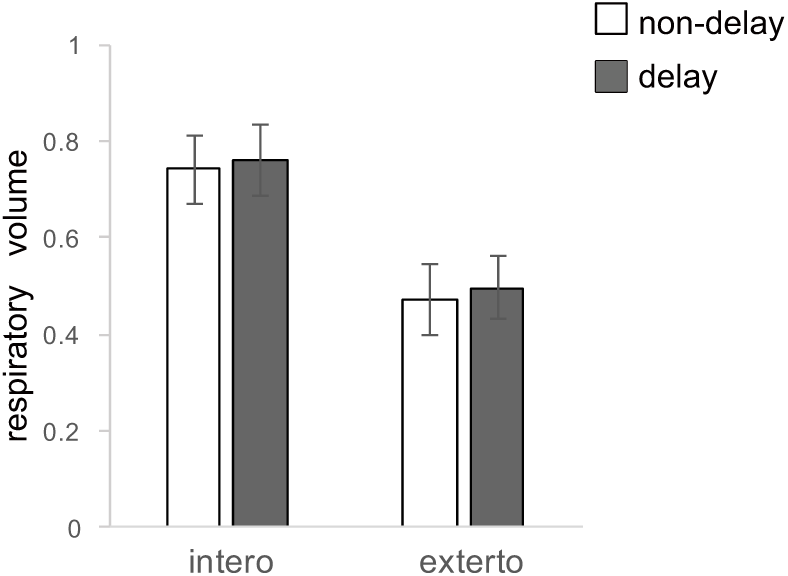
Regions showed positive (red) and negative (blue) connectivity with left AIC (as the seed) modulated by interoceptive attention (relative to exteroceptive attention) for the first fMRI sample.

**Figure 4-figure supplement 2.**
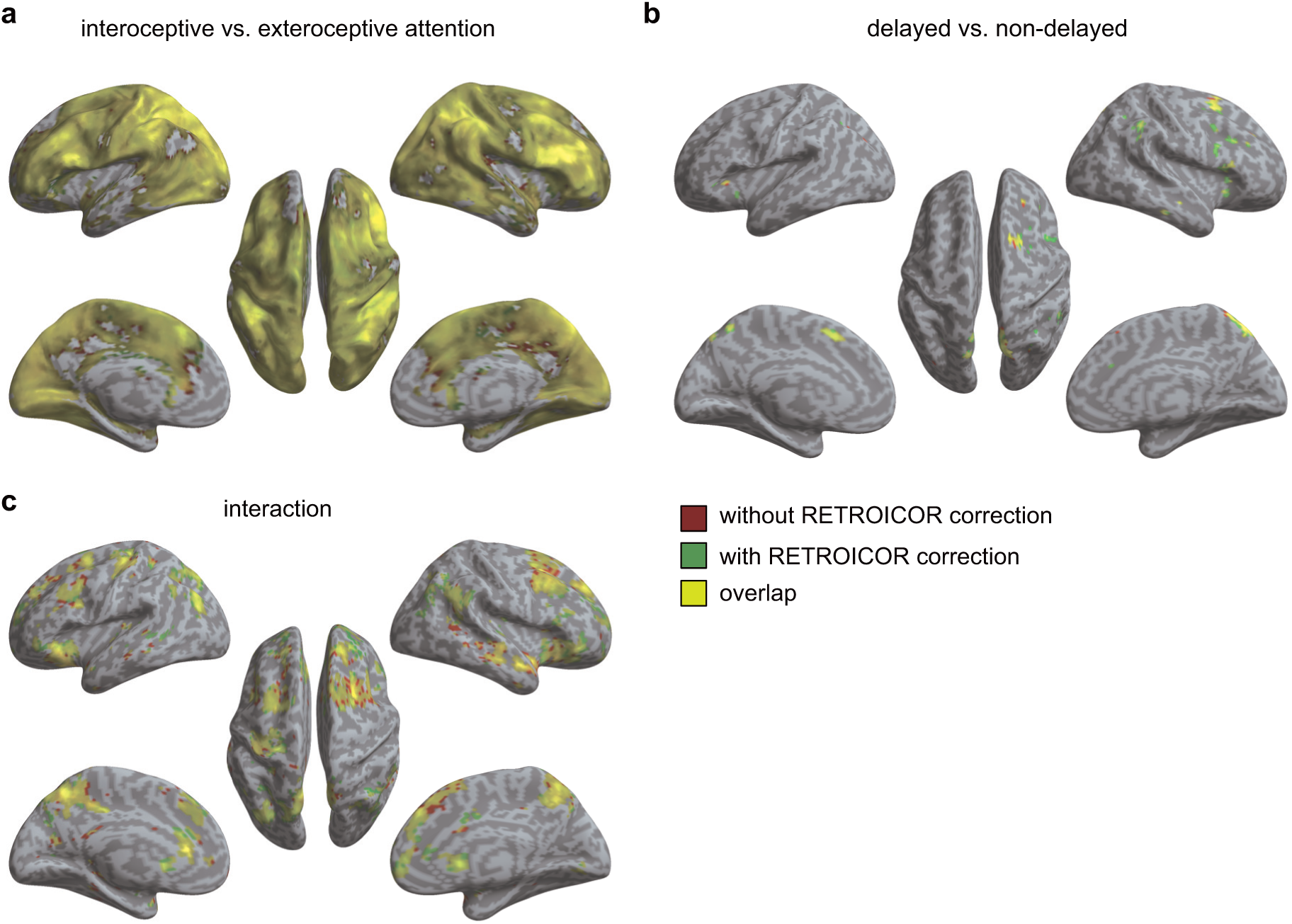
The exceedance probability of RFX BMS for the first fMRI sample. Across all 52 models, M20 outperformed the other models, and thus was identified as the optimal model. M20 denotes the model with the modulatory effects of interoceptive and exteroceptive attention exerting on the connection from the AIC to the PoCG and to V2/3.

**Figure 5-figure supplement 1.**
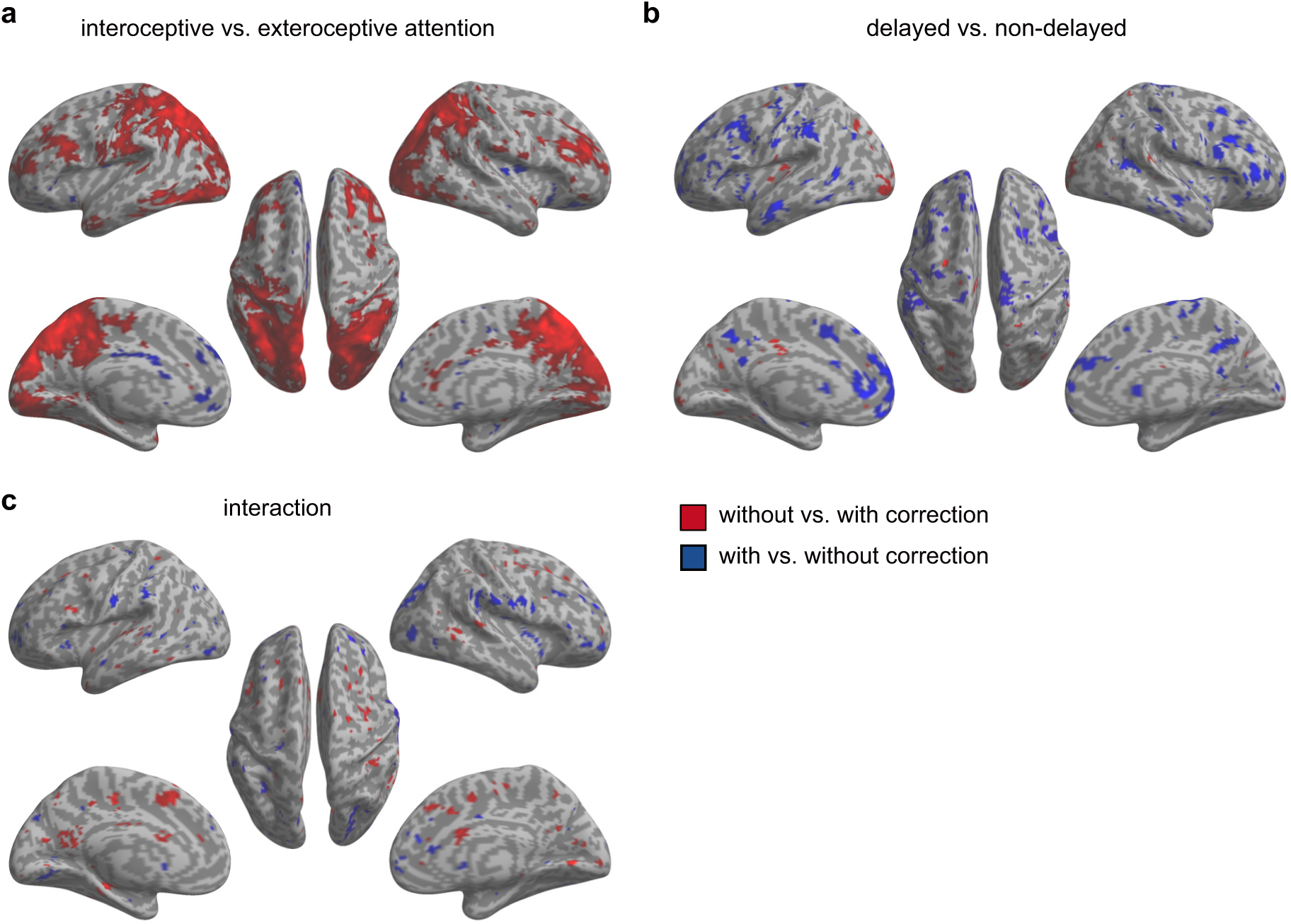
Respiratory volumes under the 4 experimental conditions from the second fMRI sample. Error bars represent 95% confidence intervals.

**Figure 5-figure supplement 2.**
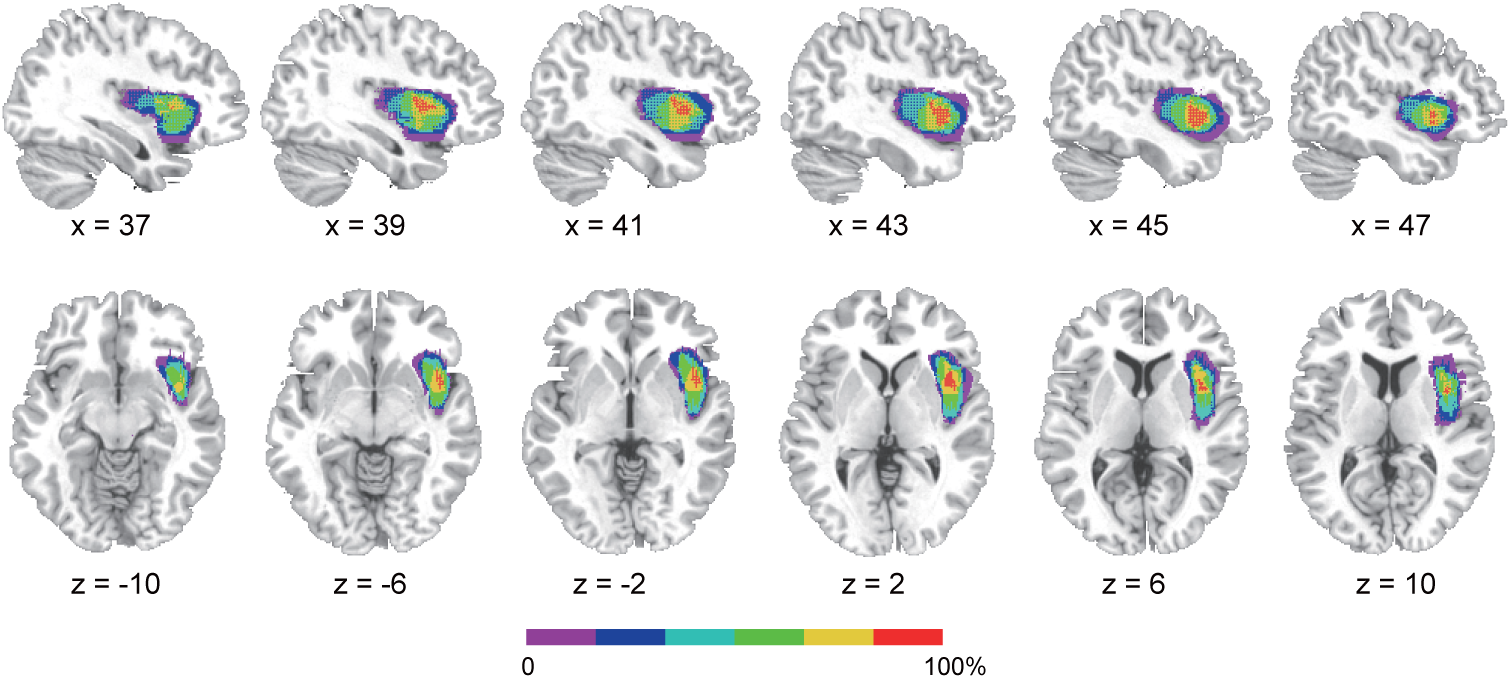
Activation maps without and with RETROICOR correction for the second fMRI sample. (a) Main effect of interoceptive attention (interoceptive task vs. exteroceptive task). (b) Main effect of breath curve feedback condition (delayed vs. non-delayed). (c) Interaction between attention type and breath-curve feedback condition ([delayed – non-delayed]_interoceptive task_ – [delayed non-delayed]_exteroceptive task_).

**Figure 5-figure supplement 3.**
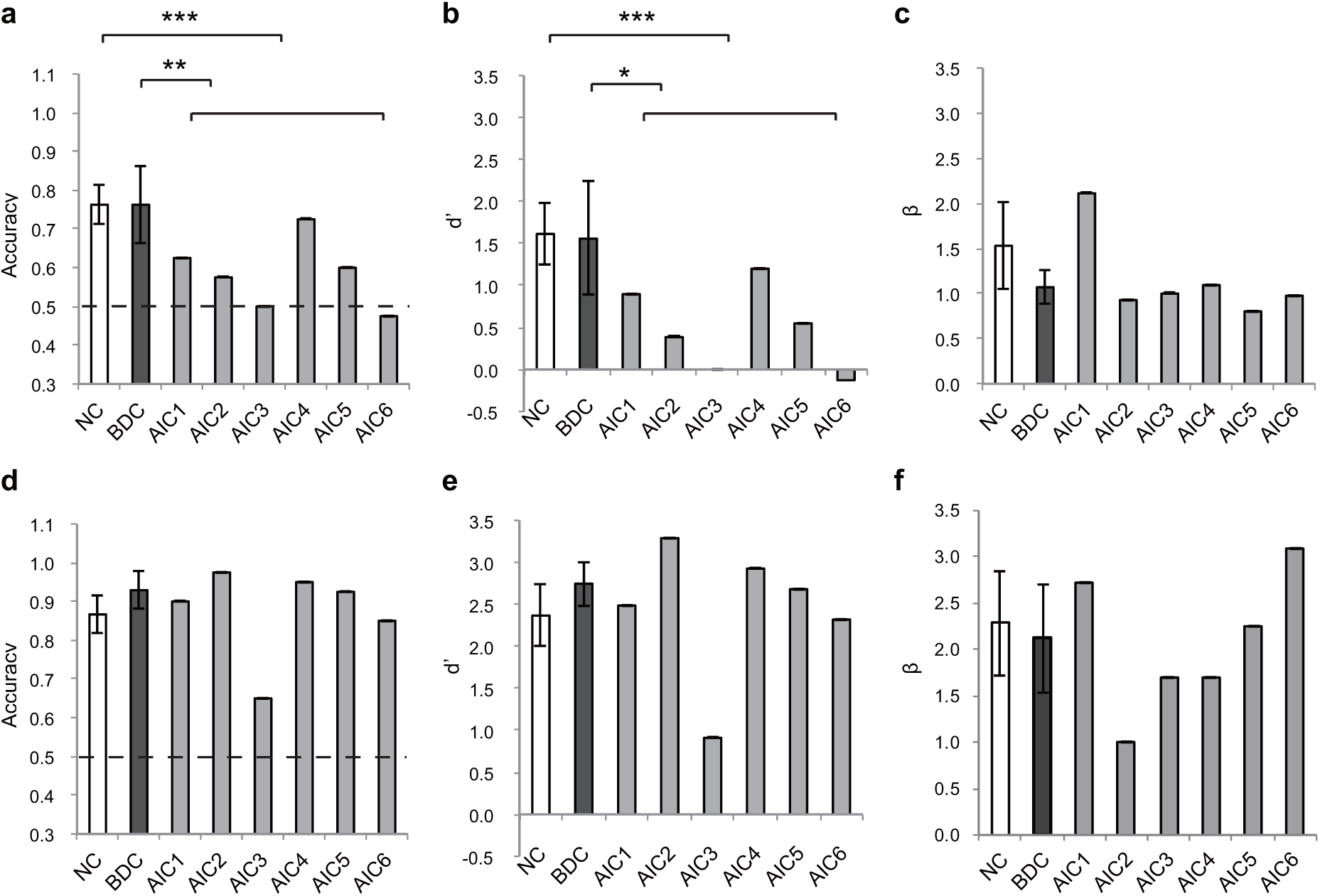
Paired t-test on beta maps obtained without and with RETROICOR correction for the second fMRI sample. The RETROICOR correction did not show a decrease on the AIC activation compared to that without the correction, suggesting that the effect of AIC was not significantly impacted by the physiological noises. (a) Main effect of interoceptive attention (interoceptive task vs. exteroceptive task). (b) Main effect of breath curve feedback condition (delayed vs. non-delayed). (c) Interaction between attention type and breath-curve feedback condition ([delayed – non-delayed]_interoceptive task_ – [delayed – non-delayed]_exteroceptive task_).

